# Combined phylogenetic and geographic data can predict plant–pest interactions with high accuracy

**DOI:** 10.1101/2024.10.30.619026

**Authors:** Elvira Hernández-Gutiérrez, Richard A. Nichols, Laura J. Kelly

## Abstract

- Non-native plant pests can pose major threats to biodiversity, with destructive ecological and economic consequences. The ability to predict future threats would allow limited resources to be concentrated on managing the most serious risks.
- We build a Bayesian model to predict hosts at risk from *Agrilus*, a beetle genus of over 3,000 species including one of the world’s worst tree pests, using phylogenetic and geographic relationships between known and potential hosts.
- We assess risk to *Quercus* (oak), their most common host, by predicting the probability of over 7,000 possible oak–*Agrilus* interactions to identify species at risk and inform future prevention efforts. Our model detects known hosts with 83.6% accuracy under Leave-One-Out cross-validation, and successfully classifies novel hosts of *Agrilus* species in new areas, indicating strong predictive performance on independent or misclassified data. Geographic proximity is a strong predictor of host sharing, with likelihood declining rapidly with distance. In general, hosts cluster phylogenetically, with a tendency for closely related oaks to share the same *Agrilus* species.
- Our approach uses readily available data and could be implemented to assess *Agrilus* interactions with other plant genera, and extended to additional host–pest systems to help prioritise counter measures against threats worldwide.

## Introduction

Throughout history, human activity has resulted in the breakdown of biogeographical barriers, prompting new biological invasions (Santini et al., 2018). In the last 200 years, the frequency of these invasions has been exacerbated by globalisation and international trade, resulting in an increase in the incidence and severity of damage by plant pests and diseases (Mack et al., 2000; Seebens et al., 2017; Brockerhoff & Liebhold, 2017; Santini et al., 2018; Seebens et al., 2025).

Although most newly introduced organisms will fail to survive, establish, and spread, a small fraction become extremely damaging, leading to cascading ecological and economic repercussions (Mack et al., 2000; Aukema et al., 2010; Brockerhoff & Liebhold, 2017). This disproportionate effect of non-native species may be explained by encounters with non-coevolved hosts that lack effective defence responses and the presence of fewer natural enemies in novel ranges (Keane & Crawley, 2002; Gandhi & Herms, 2010). Furthermore, when hosts are moved into new areas (such as through planting of tree species outside their native ranges), this can also bring them into contact with organisms with which they have not coevolved, providing further opportunity for novel interactions to arise (Pearse et al., 2013; Pearse & Altermatt, 2013).

Non-native invaders, which can remain undetected and poorly understood for decades after their introduction in a new habitat (Mack et al., 2000; Roy et al., 2014; Brockerhoff & Liebhold, 2017; Liebhold et al., 2017; Bebber et al., 2019), are difficult and costly to eradicate once established (Liebhold & Tobin, 2007). Consequently, the most efficient way to address this issue is to prevent their arrival (Mack et al., 2000). These efforts often involve trade measures such as quarantine rules (Mack et al., 2000) and phytosanitary inspections at entry points and other vulnerable locations (Wittenberg & Cock, 2001; Panzavolta et al., 2021). However, identifying threats in advance can be challenging (Pearse et al., 2013; Pearse & Altermatt, 2013) given the vast pool of organisms from which future risks could arise, and as the potential of species to cause harm often remains unrecognised until damage has already occurred (Mack et al., 2000; Liebhold et al., 2017).

Some of the most prominent forest pests and pathogens involve non-native species that have caused damage to entire ecosystems (Lowe et al., 2000; Aukema et al., 2010; Brockerhoff & Liebhold, 2017; Liebhold et al., 2023). Examples include Dutch elm disease (Karnosky, 1979), ash dieback (Enderle et al., 2019), the spongy moth (Elkinton & Liebhold, 1990; Tobin et al., 2012), and the emerald ash borer (Herms & McCullough, 2014). This last species – *Agrilus planipennis* (Coleoptera: Buprestidae) – is a beetle native to Asia, where it attacks its natural *Fraxinus* (ash) hosts only when they are already under stress (Liu et al., 2003; Wei et al., 2004). In North America, however, it has killed hundreds of millions of otherwise healthy ash trees following its accidental introduction (McCullough, 2020), causing > 99% mortality in some places (Klooster et al., 2014) and constituting the most destructive and costly forest insect invasion of the United States to date (Herms & McCullough, 2014; Fei et al., 2019).

Horizon scanning can help prevent similar outcomes to those noted above by narrowing the pool of taxa on which surveillance and mitigation efforts should focus, for example, by highlighting organisms that warrant targeted monitoring or restrictions on the movement of high-risk materials (DEFRA, 2025; Peyton et al., 2025; Ribaya et al., 2025). This can, however, be a time-consuming and labour-intensive process (Antoniou et al., 2024), often requiring detailed evaluation of individual species (e.g., see the UK Plant Health Risk Register, DEFRA, 2025). Furthermore, these assessments tend to focus heavily on organisms with already documented impacts, and are therefore likely to overlook novel threats (Hulme, 2025). Our intention is to address this challenge by providing a modelling-based workflow that predicts highly likely plant–pest interactions, efficiently identifying pest candidates for in-depth assessment and helping to direct resources towards those most likely to pose a threat.

Previous research has shown that there is often a phylogenetic signal in the host ranges of plant enemies, meaning that more closely related plants tend to share the same pests or pathogens (Parker & Gilbert, 2004; Pearse & Hipp, 2009; Yguel et al., 2011; Gilbert & Parker, 2016). Consequently, models informed by phylogenetic relationships between known and potential hosts have been developed in efforts to predict novel biotic interactions to facilitate the assessment of risks to plant health (e.g., Gilbert & Parker, 2016; Robles-Fernández & Lira-Noriega, 2017). However, this single predictor may not always be sufficient to provide accurate results, particularly at finer phylogenetic scales (Cavender-Bares et al., 2006) due, for example, to phenotypic convergence reducing the strength of signal (Becerra, 1997; Pearse et al., 2013). Therefore, models integrating additional variables, such as geographic, trait, and environmental data, have also been used to predict novel interactions in a range of taxonomic groups (Novotny et al., 2010; Gilbert et al., 2012; Pearse & Altermatt, 2013; Albery et al., 2020; Farrell et al., 2022; Gougherty et al., 2024), or to estimate the probability of infection or damage from known pests and pathogens (Barrow et al., 2019; Mech et al., 2019). However, approaches incorporating multiple predictors can pose challenges in scalability and complexity, and information such as trait data can be incomplete or hard to compile (Albery et al., 2020). For example, Mech et al. (2019) excluded certain traits from their modelling of risks from herbivorous insects where data were unavailable for the majority of host species, and Barrow et al. (2019) had to infer trait status from related taxa in some cases when modelling parasite infections in birds. Moreover, Barrow et al. (2019) found species-level predictors of host susceptibility, including environmental variables such as temperature and aridity, differed in importance depending on the genus of parasites under consideration. These examples emphasize the challenge of identifying predictors to supplement phylogenetic signal that can be broadly applied.

Here, we develop a statistical model to predict potential novel host plant associations of *Agrilus*, in order to anticipate future threats from this major pest-containing genus. Our aim is to develop a tractable approach that relies only on information that is readily available for many plant taxa – host phylogenetic relatedness and geographic overlap. We hypothesise that incorporating geographic overlap between known and potential novel hosts of *Agrilus* beetles into phylogenetically informed statistical models will improve their power to predict novel interactions because host jumps are more likely between plants occurring in closer physical proximity (Parker & Gilbert, 2004). Although host geographic overlap has been found to be predictive of pest and pathogen sharing between mammal species (Albery et al., 2020; Brennan et al., 2024), to the best of our knowledge, it has not been used used to aid prediction of novel interactions between plants and their pests.

We use *Agrilus* as our test case since multiple *Agrilus* species, including the emerald ash borer, are of significant concern to plant health (Nielsen et al., 2011; Herms & McCullough, 2014; Lopez et al., 2014; Volkovitsh et al., 2020). With more than 3,000 described species worldwide (Jendek & Grebennikov, 2023), the number of potential pest species that could emerge from within this genus alone is substantial. We focus on *Agrilus* species that exploit oaks (*Quercus* spp.), their most common host genus (Jendek & Poláková, 2014), and one of major ecological, cultural, and economic importance (Tantray et al., 2017). We test for phylogenetic signal in *Agrilus* host use, and evaluate whether combining phylogenetic relatedness with geographic proximity yields accurate predictions for *Quercus–Agrilus* interactions. Our approach provides a straightforward framework for identifying potential novel hosts of *Agrilus* species and offers a starting point for targeted early-detection efforts and for informing policies designed to reduce the risk to plant health. Moreover, it can be used for predicting novel interactions both at the establishment phase in the scenario of pests invading new areas, and where potential hosts have been introduced into the range of pests. While developed for *Agrilus*, our framework could be applied in other systems where trait data are sparse but phylogenetic and geographic information is accessible, allowing large numbers of species to be analysed simultaneously and enabling identification and prioritisation of potential pests for more in-depth assessment.

## Materials and Methods

See **Fig. 1** for an overview of the methods.

**Fig. 1:**
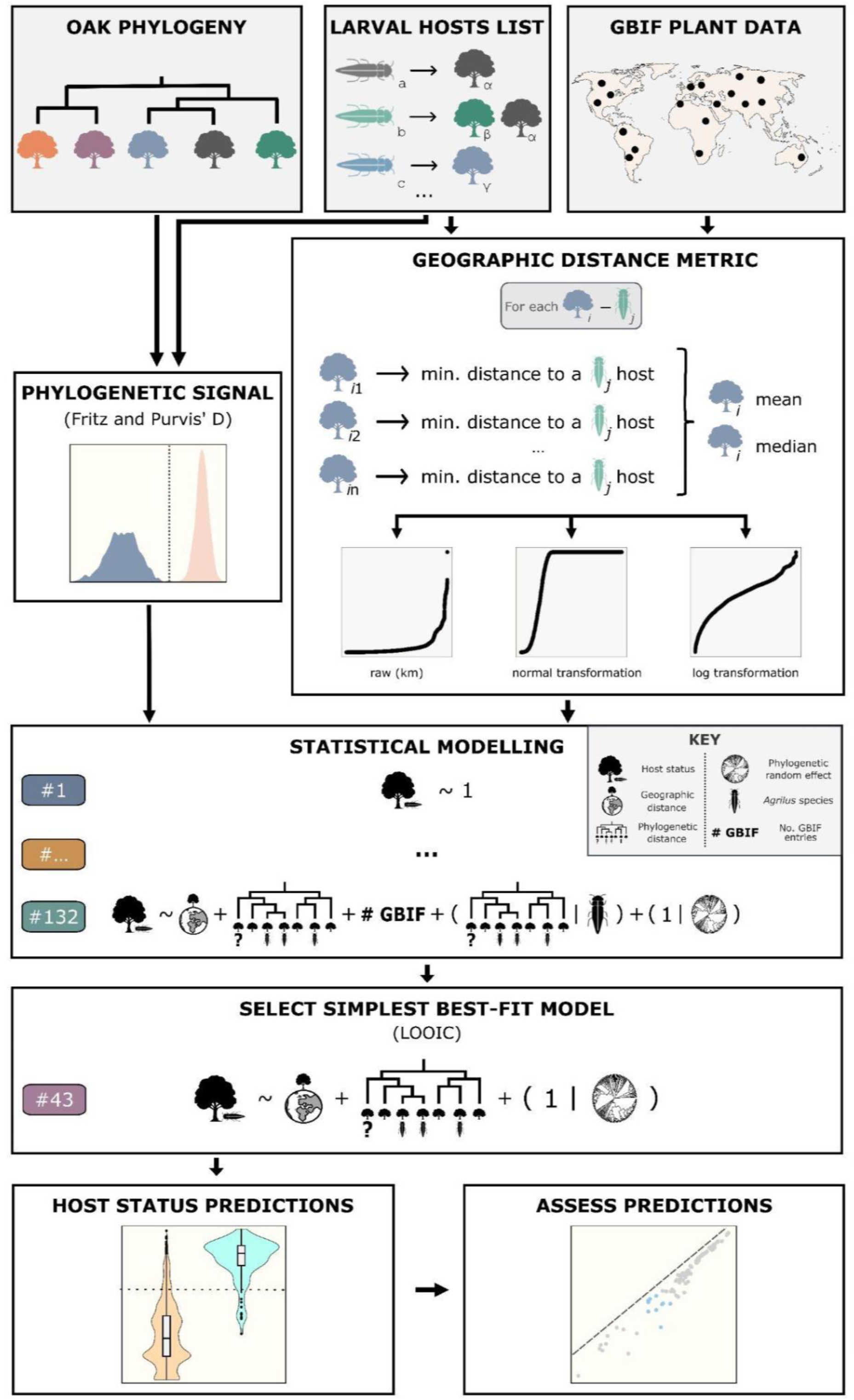
Schematic overview of the study pipeline. We retrieved a *Quercus* phylogenetic tree (Hipp et al., 2020), produced a list of larval hosts of *Agrilus* beetles that utilise oaks (Vansteenkiste et al., 2004; Nielsen et al., 2011; Brown et al., 2015; Cipollini, 2015; Jendek, 2016; Peterson & Cipollini, 2017; Prokhorov & Vasilyeva, 2017; Jendek & Nakládal, 2019a, 2019b, 2019c; Volkovitsh et al., 2020) and downloaded plant geolocation information from the Global Biodiversity Information Facility (GBIF, 2022). We cleaned the data and then tested for the presence of a phylogenetic signal in our data using Fritz and Purvis’ D statistic (Fritz & Purvis, 2010). Next, we computed several geographic distance metrics, compared them and subsequently built multiple binary logistic models (Bürkner, 2017) using different combinations of several explanatory variables to assess which have a significant effect on *Agrilus* host status. We employed the Leave-One-Out cross-validation Information Criterion (LOOIC) (Vehtari et al., 2017) for model selection, and used the chosen model to predict *Agrilus* host status for all *Quercus* species – *Agrilus* species pairs. Finally, we performed Leave-One-Out (LOO) analyses to assess the ability of our model to correctly identify known hosts.

### Assembly of data

Information on worldwide *Agrilus* Curtis, 1825 host plants was extracted from an extensive critical review of the literature (Jendek & Poláková, 2014). These data were supplemented by records from subsequent publications (Vansteenkiste et al., 2004; Nielsen et al., 2011; Brown et al., 2015; Cipollini, 2015; Jendek, 2016; Peterson & Cipollini, 2017; Prokhorov & Vasilyeva, 2017; Jendek & Nakládal, 2019a, 2019b, 2019c; Volkovitsh et al., 2020) and formatted as a dataframe, with information available for a total of 241 *Agrilus* species. Plants were considered hosts only if there were records of beetle larvae exploiting them, as (i) larval host data is more reliable (since adults could be present on trees they do not exploit), and (ii) it is the larvae that inflict significant host damage. For records from Jendek & Poláková (2014), we only retained those with a confidence index of 3 (highest confidence), at least one larval reference (marked as “!”), and no remarks indicating the data might be unreliable. All plant taxon names were standardised against the World Checklist of Vascular Plants (WCVP) v3 (Govaerts et al., 2021). For subsequent analyses, this dataset was restricted to *Agrilus* species that utilise *Quercus* L., retaining all confirmed larval host species reported within their native ranges, including non-native hosts (**Table S1**). In the case of *A. auroguttatus*, *A. bilineatus* and *A. sulcicollis,* all of which have been introduced outside their native range, any non-native hosts only recorded in these new areas were removed. Our final dataset contained information for 32 *Agrilus* species that are known to use, in total, 51 *Quercus* species. Continent-level native distribution data in **Table S1** were inferred from Jendek & Grebennikov (2023) and GBIF (2025) for *Agrilus* species, and from WCVP v10 distribution data (Govaerts et al., 2021) for host taxa.

A time-calibrated phylogenetic tree for *Quercus* (Hipp et al., 2020) was retrieved from https://github.com/andrew-hipp/global-oaks-2019 (file tr.singletons.correlated.1.taxaGrepCrown.tre). It represents ca. 250 *Quercus* species (> 50% of all recognised oak species in WCVP v3), and was inferred using restriction-site associated DNA sequencing and with fossil calibration (Hipp et al., 2020); we chose to use this tree as it was the most comprehensive time-calibrated species-level phylogenetic tree for *Quercus* available at the time of our analyses. Although Hipp et al. (2020) also included an equivalent tree using stem calibration, phylogenetically informed models of pest host use are robust to the precise choice of host phylogenetic tree (Pearse & Hipp, 2014; Uden et al., 2022). Species names in this tree were standardised against the WCVP v3; intraspecific taxa were reduced to species-level, and one tip per species retained. One known *Agrilus Quercus* host (*Q. gambelii*) is missing from this phylogenetic tree and was therefore removed from our dataset. The final modified tree contained 238 *Quercus* species, including two new, uncharacterised species, which were retained for the phylogenetic signal testing but excluded from the statistical modelling analyses as they lack geographic information.

Global geographic distribution records for *Quercus* species and any other known host taxa of the 32 *Agrilus* species was obtained from the Global Biodiversity Information Facility on June 19th, 2022 (GBIF, 2022), using the rgbif R package v3.7.3 (Chamberlain et al., 2003). This dataset was cleaned, following recommendations from the CoordinateCleaner R documentation (Zizka et al., 2019): we removed entries with unreliable coordinate information, flagged as “COORDINATE_INVALID”, “COORDINATE_OUT_OF_RANGE”, “COORDINATE_REPROJECTION_SUSPICIOUS”, “COORDINATE_UNCERTAINTY_METERS_INVALID”, “COUNTRY_COORDINATE_MISMATCH”, “PRESUMED_SWAPPED_COORDINATE”, “PRESUMED_NEGATED_LONGITUDE”, “PRESUMED_NEGATED_LATITUDE”, or “ZERO_COORDINATE”. We also excluded records with unusually high individual counts (> 10), from fossils, or predating 1950, since these may less accurately reflect current species occurrences (see “Improving data quality” recommendations in https://cran.r-project.org/web/packages/CoordinateCleaner/vignettes/Cleaning_GBIF_data_with_CoordinateCleaner.html, Zizka et al., 2019). The set was further filtered using the clean_coordinates() function from the CoordinateCleaner R package v2.0-0 (Zizka et al., 2019) (testing for “capitals”, “centroids”, “duplicates”, “equal”, “outliers”, “gbif”, “institutions”, “zeros” and “seas”, and with outliers_method = “distance”, capitals_rad = 1000, inst_rad = 1000, outliers_size = 50, zeros_rad = 1 and all other parameters set default). Due to memory constraints, species with > 100,000 georeferenced entries following cleaning were randomly subsampled to 100,000. Finally, a single manual coordinate entry was added for oak species in the phylogeny lacking georeferenced information (i.e.*, Q. sagrana* and *Q. yiwuensis*); their native ranges were checked via https://powo.science.kew.org, and the coordinates for their region centroid according to CoordinateCleaner extracted.

### Phylogenetic signal

We tested for the presence of a phylogenetic signal in host range using Fritz and Purvis’ D statistic (Fritz & Purvis, 2010) – which can be used for binary traits – implemented in the R package caper v1.0.1 (Orme et al., 2023), using 1,000 permutations. To visualise this signal and verify the pattern, we looked for clades in the oak phylogeny with an over-representation of *Agrilus* hosts (host-rich clades, **Fig. 2**), using an approach originally developed for analyses of community structure (C. O. Webb et al., 2008) but since used to identify clades with overrepresented traits (see Saslis-Lagoudakis et al., 2012, and Holzmeyer et al., 2020). To identify host-rich clades, we used the nodesig function from the phylocom v4.2 software (C. O. Webb et al., 2008), with one million randomisations and the default swap method (method 2). The input files were the *Quercus* phylogenetic tree (file tr.singletons.correlated.1.taxaGrepCrown.tre, noted above, but modified to remove bootstrap support values), a traits file with the type of trait set to binary (type 0), and a sample file with the first column indicating host status and all abundances set at 1.

**Fig. 2:**
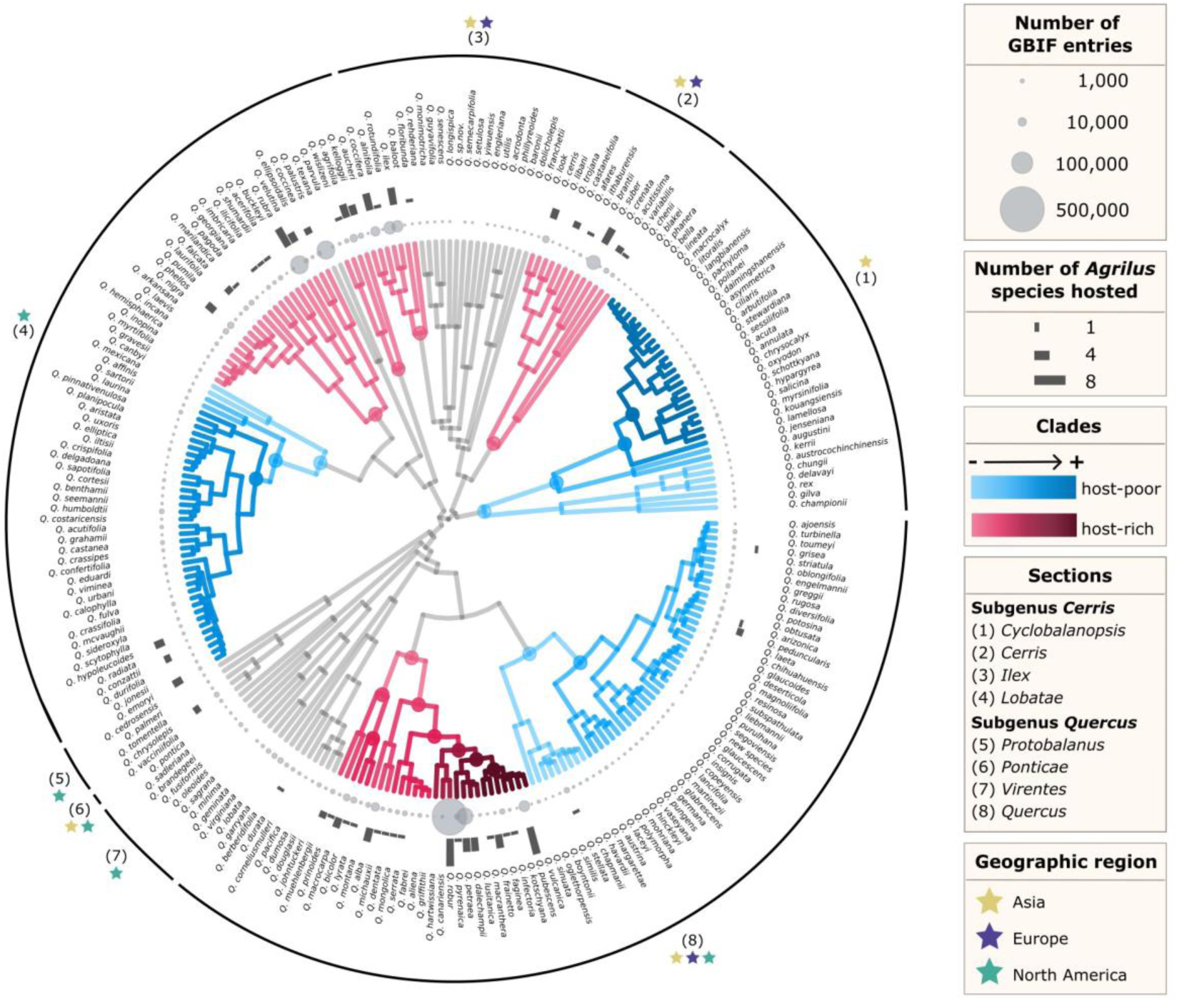
Phylogenetic patterns of clustering of the *Agrilus* hosts across the *Quercus* phylogeny. Red and blue dots indicate nodes identified as having significantly more or less descendants that are hosts than expected by chance, respectively. Grey clades indicate no over-or under-representation of hosts. We identified five main clades with an over-representation (i.e., a significant degree of clustering) of *Agrilus* hosts (host-rich clades, in red). One of these clades represented seven additional nested host-rich clades (in section *Quercus*). Non-hosts present in these clades could be potential future hosts at risk. There are also three host-poor clades (in blue) where hosts are underrepresented. Non-hosts present in these clades may be at a lower risk of future host jumps by *Agrilus*. However, host-poor clades for *Agrilus* as a genus might still be host-rich for particular *Agrilus* species (for example, *A. auroguttatus* utilises three *Quercus* species present in three nested host-poor clades). Dark grey barplots indicate the number of known *Agrilus* species hosted per *Quercus* species, and light grey circles indicate the number of cleaned GBIF entries (see **Materials and Methods**) available for each. Stars indicate geographic regions for major *Quercus* clades according to (Kremer & Hipp, 2020), and numbers indicate sections (see key). It should be noted that this clustering analysis is scale dependent.

### Statistical modelling

We built generalised linear mixed-effects models (or multilevel models, in the terminology of the brms R package, Bürkner, 2017) using binary logistic regressions to assess whether host status (i.e., a binary trait indicating whether an oak species hosts a given *Agrilus* species, where 0s indicate oak species not recorded as a larval host of that particular *Agrilus* species in our dataset, and 1s indicate oak species that are confirmed larval hosts; see **Assembly of data**) can be explained by the *Quercus* phylogeny and the geographic proximity to known *Agrilus* host plants.

Since no standard measure of geographic proximity between species exists for these purposes, we evaluated six metrics of spatial overlap between oak species and *Agrilus* hosts (see **Notes S1** for their functional definitions), calculated from the GBIF geolocation records (GBIF, 2022). For our first two metrics, for each possible *Quercus* species–*Agrilus* species pair, we calculated the mean and median of the minimum distances (in km) between each record of the given oak species and any known plant host (i.e. including non-*Quercus* hosts) of the given *Agrilus* species. If the focal *Quercus* species was a known host, the distance was computed to any *other* host species, and if it was the only known host, it was set to a large value of 30,000 km to represent the absence of alternative hosts. Distances were calculated using the geodist() function from the R package geodist v0.0.7 (Padgham M, 2021), with “measure” set to “cheap” due to time and memory constraints. For the next two metrics, we weighted distances within a plausible dispersal range (C. R. Webb et al., 2021; Schrader et al., 2020) most heavily. Specifically, we transformed the mean or the median of the minimum distances to the corresponding normal probability density (μ = 0 km, σ = 50 km), such that shorter distances received higher weights. We then multiplied this value by −1, so that species close in space had lower scores, and rescaled the resulting value to range from 0 to 1 for ease of interpretation. For the final two metrics, the original distances were instead log-transformed and then rescaled to between 0 and 1. We then built separate models using each of these six distance metrics in turn (see **Notes S2**) and compared them using a Leave-One-Out (LOO) analysis (Vehtari et al., 2017), using the loo() function from the brms v2.18.0 package, with moment_match = TRUE and reloo = TRUE to deal with high Pareto *k* diagnostic values (we note that only two out of 7552 observations had a high diagnostic value). The two normal-transformed distances consistently outperformed the others (**Table S2**) and were selected as the best-performing metrics for use in the subsequent analyses.

Additionally, we computed three phylogenetic metrics for each *Quercus*–*Agrilus* interaction using the time-calibrated *Quercus* phylogenetic tree (see **Notes S3** for their functional definitions). These were (i) the mean phylogenetic distance (where the distance to the root = 0.5) of the given oak species to (other) *Quercus* hosts of the given *Agrilus* beetle, capturing the overall phylogenetic breadth of its known hosts; (ii) the minimum phylogenetic distance of the given oak species to any (other) hosts, capturing the proximity to the closest known host; or (iii) the sum of both. If the current oak species was the only known host, distances were set to twice the distance to the root.

We then fitted binary logistic models with brms (Bürkner, 2017), using the brms() function with family = “bernoulli”, iter = 7000, cores = 4, control = list(adapt_delta = 0.995, max_treedepth = 15), and default settings for all other parameters. We modelled host status as a binary response (0 for non-hosts, 1 for hosts) and compared 132 models comprising all possible combinations of explanatory variables that included: (i) one of the two best-performing geographic overlap metrics (fixed effect), (ii) one of the three computed specific phylogenetic metrics (fixed effect), (iii) a proxy for sampling effort (fixed effect), i.e., number GBIF records per oak species, as GBIF data are known to reflect this bias (Beck et al., 2014; Maldonado et al., 2015), (iv) a general *Quercus* phylogenetic random effect (with one level per oak species, so that oaks closely related to species that host *Agrilus* have a greater chance of being hosts) and (v) an *Agrilus* species random effect. The simplest model was an intercept-only null model (**Table S3**). For those models with a phylogenetic random effect, this was included as a varying intercept over oak species, which were correlated as specified by a phylogenetic covariance matrix (see https://cran.r-project.org/web/packages/brms/vignettes/brms_phylogenetics.html, Bürkner, 2017). To test whether the influence of host relatedness varies among *Agrilus* species, models including both a fixed phylogenetic variable and a random *Agrilus* species effect were fitted in two ways (**Table S3**): (i) keeping the original *Agrilus* species random effect (intercept-only), and (ii) transforming it into a random-slope model to allow the strength of the phylogenetic effect to vary among *Agrilus* species. This reflects the hypothesis that generalist species are less constrained by host relatedness than specialists.

To identify the best-fit models, we employed a model comparison approach using Leave-One-Out Cross-Validation Information Criterion (LOOIC; Vehtari et al., 2017), using the brms loo() function as specified above. Due to memory limitations, the models were divided into three groups for initial comparison and the best-performing models from each group were then compared against each other to identify the best-performing model overall (**Tables S4–12**). This final model was:

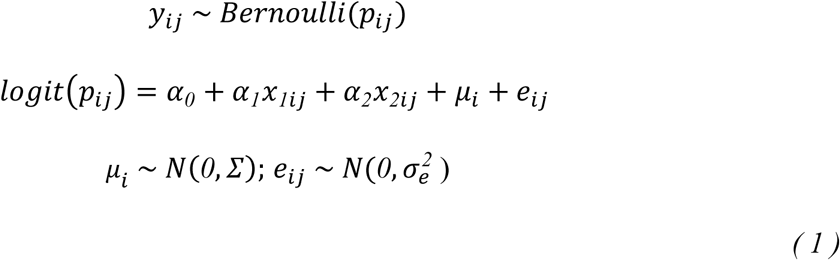

𝑦_𝑖𝑗_𝜖 {*0*,*1*} indicates if *Quercus* species 𝑖 can host the *Agrilus* species 𝑗. The probability 𝑝_𝑖𝑗_ is a function of the following variables. The value 𝛼*_0_* is the intercept, 𝑥*_1_*_𝑖𝑗_ is the normal-transformation of the mean minimum geographic distance from the *Quercus* species 𝑖 to any (other) known host of *Agrilus* species 𝑗, and 𝛼*_1_* is the associated coefficient, 𝑥*_2_*_𝑖𝑗_ is the minimum phylogenetic distance of the *Quercus* species 𝑖 to any (other) known oak host of the *Agrilus* species 𝑗, and 𝛼*_2_* is the associated coefficient, *µ*_𝑖_ is the phylogenetic random effect with its covariance matrix 𝛴 specified by the *Quercus* phylogeny and an additional parameter 𝜎*^2^*, and 𝑒_𝑖𝑗_ is the normally distributed residual error.

### Predicting host–*Agrilus* interactions

We used the model in **Equation (1)** to compute predictions for the response variable (i.e., host status for each *Quercus* species – *Agrilus* species pair) with the brms fitted() function (using scale = “linear” to maintain the log-odds scale). To classify which *Quercus* species might be particularly at risk, we also computed a binary threshold to divide predictions into 0s (i.e., non-hosts) and 1s (i.e., hosts). This cut-off was selected as the point that maximises sensitivity and specificity (i.e., where their functions intercept, see **Fig. S1**).

To evaluate the model performance, we compared two data limitation scenarios : one in which we falsely assume there is no interaction (misclassified data), the second in which no information is available for the given interaction (cross-validation using independent data). For the first, we ran a deliberately conservative analysis where, in turn, each known host–*Agrilus* interaction (i.e., coded as 1 in the original model) was intentionally misclassified as a 0, and the value for this observation predicted (using fitted() as above). For the second case, we incorporated the absence of knowledge about the interaction using the brms loo_linpred() function (with type = “mean”), which draws from the leave-one-out distribution using Pareto-Smoothed Importance Sampling. This procedure uses the model’s posterior distribution as the proposal distribution for importance sampling (Vehtari et al., 2017). In addition, since k-fold cross-validation is a commonly used approach for testing model performance, we also compared the predictions of the second LOO method against those obtained from five-fold cross-validation. The five-fold analysis was performed using brms functions kfold() (using save_fits = TRUE) and kfold_predict() (using method = “posterior_epred”). The predictions for the missing observations were then averaged across posterior draws and transformed to the log-odds scale for comparison.

All analyses were performed in R v4.2.0.

## Results

Our final dataset includes 32 *Agrilus* species from Africa, Asia, Europe, North and South America, with 40.63% (13) of species recorded from a single continent and 46.88% (15) found in all five (**Table S1**). Overall, 56.25% (18) of these *Agrilus* species have *Quercus* as their only reported hosts (**Table S1**). Host breadth averages 4.53 plant species (range = 1–15) and 2.03 genera (range = 1–9) per *Agrilus* species (**Table S1**). The average number of hosted *Agrilus* species for the 50 *Quercus* hosts present in the phylogenetic tree is 2.32 (range = 1–8; **Fig. 2** and **Table S1**).

### Phylogenetic signal in host status

*Agrilus* beetle hosts are non-randomly distributed across the *Quercus* phylogeny (Hipp et al., 2020), with several clades showing strong clustering according to the phylocom analysis (**Fig. 2**). On average, for each *Agrilus* species there are 3.63 reported *Quercus* host species (range = 1–10), and 232.38 non-host *Quercus* species (range= 226–235) in our dataset. Hosts are over-represented in section *Cerris* (with the entire section identified as a host-rich clade), as well as in one clade of section *Ilex* and two of section *Lobatae* (**Fig. 2**). Clades where *Agrilus* hosts are under-represented (host-poor clades) are found within section *Cyclobalanopsis*, section *Quercus* and interestingly, given the presence of two host-rich clades, also within section *Lobatae* (**Fig. 2**). These findings are consistent with the presence of a moderately strong overall phylogenetic signal in host status indicated by Fritz and Purvis’ D (D = 0.56 Fritz & Purvis, 2010; **Fig. 3a**), where the distribution differs significantly from both a Brownian expectation (p = 0.001) and a random one (p < 2e-16; **Fig. 3a**).

**Fig. 3:**
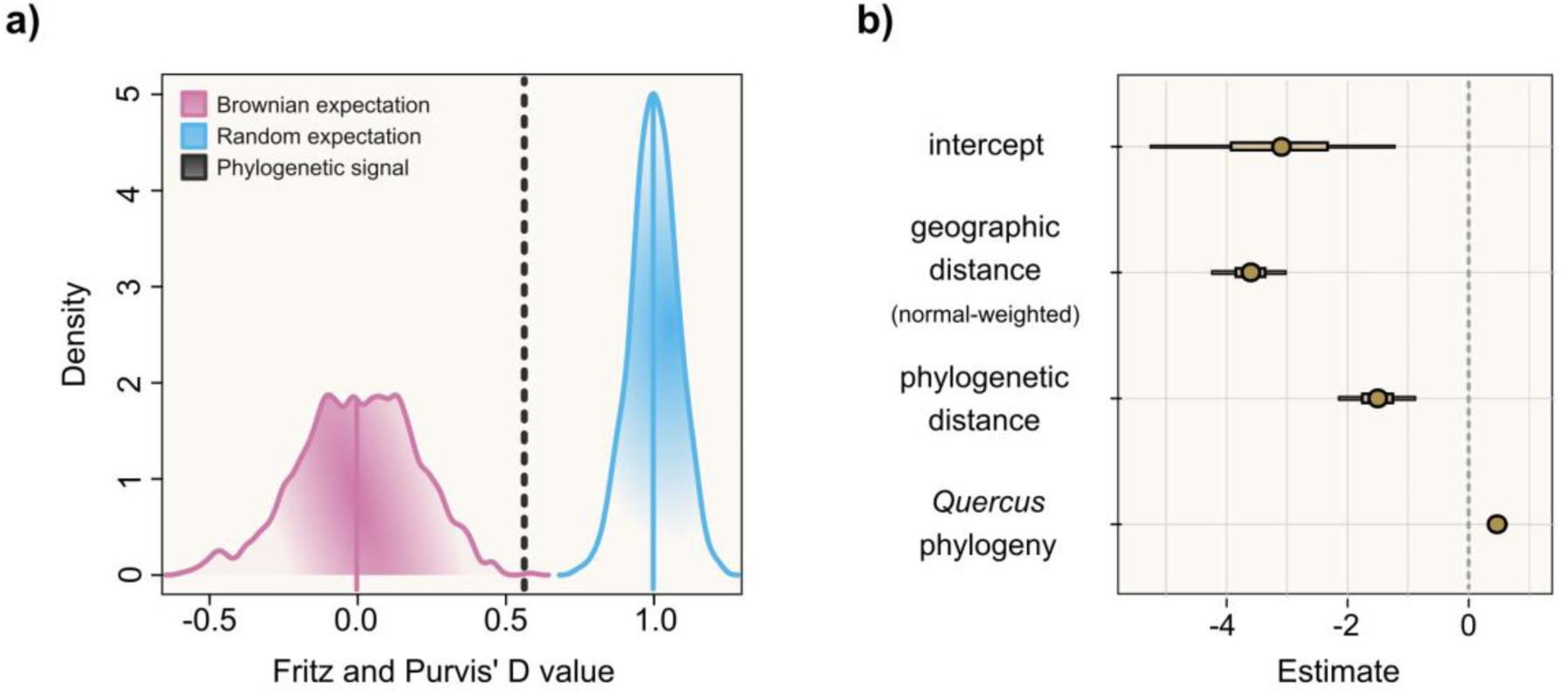
Phylogenetic signal, and posterior mean estimates and 95% Credible Intervals (CI) for the chosen model, i.e., model 43. **a) Fritz and Purvis’ D statistic for the distribution of *Agrilus* hosts in the *Quercus* phylogeny.** We found significant levels of clumping of *Agrilus* hosts, with the phylogenetic signal being moderately strong (D = 0.56, dotted line). Our signal differs significantly from (i.e., does not overlap with) both the Brownian (pink distribution) and the random expectation (blue). **b) Posterior mean estimates and 95% Credible Intervals (CI) of predictors and random effects on host status for model 43.** In this binomial model, related *Quercus* (oak) species have similar probability of hosting *Agrilus*, i.e., the variation among the 𝜇_𝑖_ (phylogenetic random effect) is phylogenetically correlated (intercept σ = 0.48, CI = 0.31 – 0.69). This means that species have different baseline probabilities of being hosts, with more closely related species having more similar values. See **Table S13** for the group-level (i.e., random) effects of each level (*Quercus* species). The estimates of *α_2_* (coefficient of the species-specific phylogenetic effect) show that closely-related oak species are more likely to be hosts of a particular *Agrilus* species (estimate = −1.51, 95% CI = −2.30 – −0.75). The estimates of *α_1_* (coefficient of the geographic distance effect) show that oak species spatially close to hosts of a given *Agrilus* species, *j*, are more likely to also be hosts (estimate = −3.61, CI = −4.39 – −2.90). Given that all CIs are narrow, the effect of these variables appears to be strong, albeit with the effect of the (fixed) phylogenetic distance being the least pronounced (**Fig. S2**). The negative overall intercept (−3.16, CI = −5.82 – −0.93) indicates that the naive probability of any given random *Quercus* species being a host is low. This model did not perform significantly worse than those that included a random effect for *Agrilus* species or a proxy for sampling effort, indicating that this simpler model can be applied to predict hosts for *Agrilus* species with limited available information.

### Variable predictors of *Agrilus* host status

Among the 132 candidate models (see **Table 1** for a summary and **Tables S4–S12** for full statistics and comparisons), our selected model was the one best supported by the LOOIC comparisons (**Table S8**; see **Materials and Methods**). Only one other best-performing model (model 44, **Table 1**) is equally simple; the remaining eleven models are more complex (**Tables 1 and S8**).

**Table 1:**
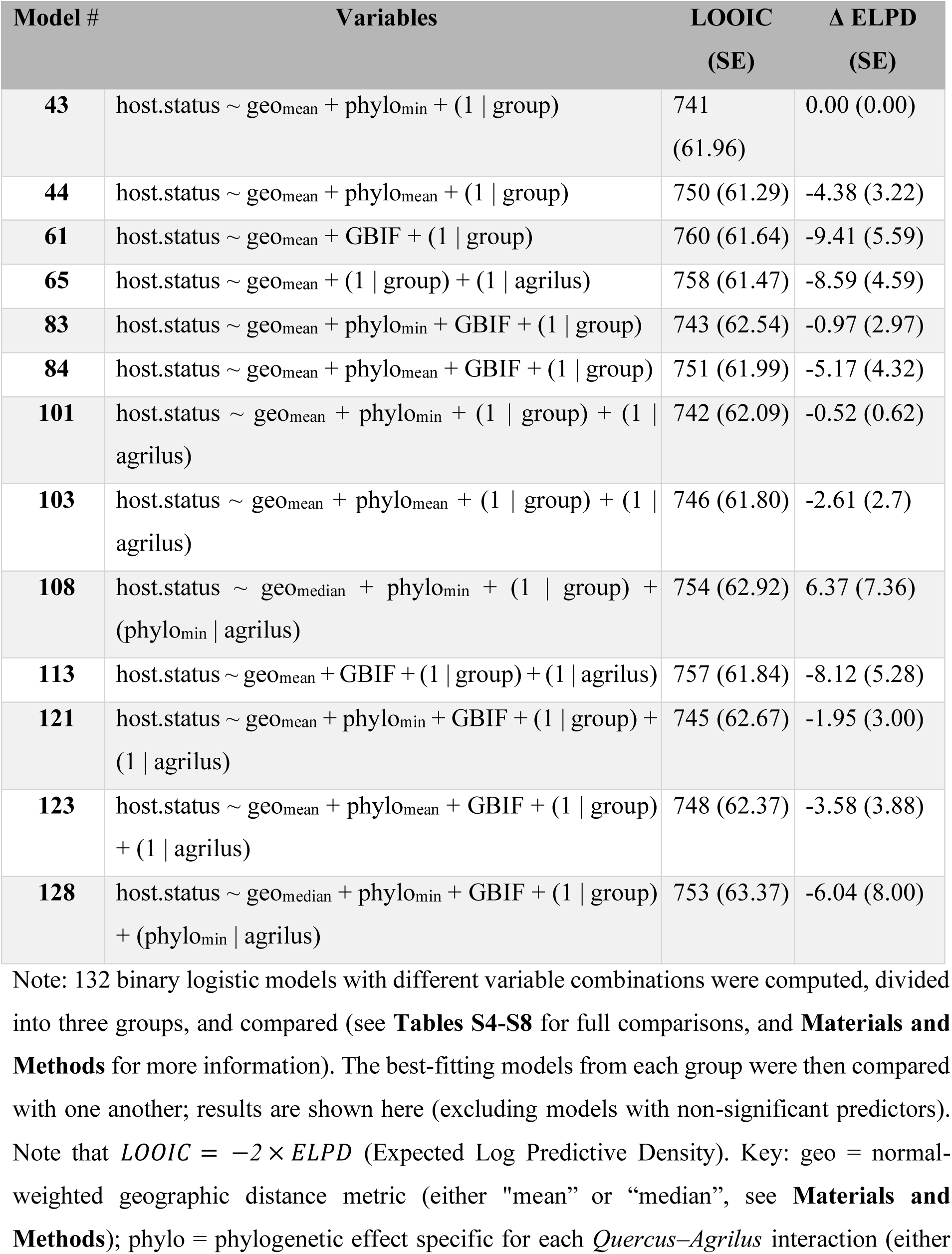
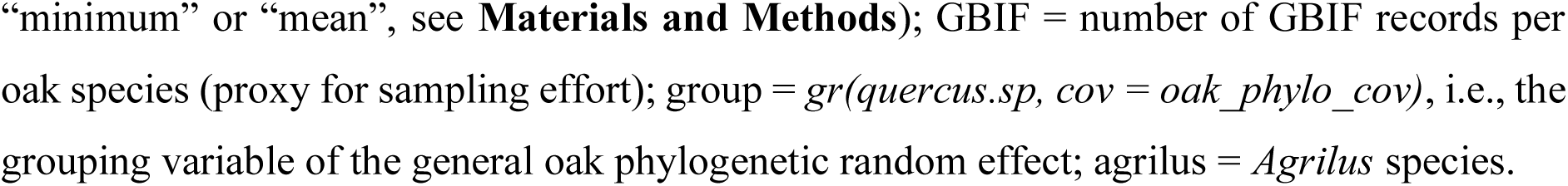
Comparison of selected computed statistical models explaining *Agrilus* host status of *Quercus* species.

Our final model, **Equation (1)**, reveals that *Agrilus* beetle host use in *Quercus* is governed by both phylogenetic and spatial structure (**Fig 3b**). Host sharing is strongly influenced by geographic proximity (**Fig. 3** and **Fig. S2**): oak species are more likely to host a given *Agrilus* species (*j*) if they grow near another host. This spatial effect, quantified by *α_1_*, declines rapidly with distance, with the best-performing metric for 𝑥*_1_*_𝑖𝑗_ being the normal-transformed distance (σ = 50 km). Model comparison using LOOIC confirms the importance of this effect, as models including *α₁* consistently outperform those without it, i.e., with *α_1_* = 0 (e.g., model 2 vs. 1 and 121 vs. 115; **Tables S9 and S11**).

The model also captures two distinct phylogenetic effects on host status. First, there is a strong general tendency for *Agrilus* hosts, regardless of which *Agrilus* species they host, to be phylogenetically clustered (**Fig. 3** and **Fig. S2**). This is reflected in the phylogenetic correlation of the variation among the 𝜇_𝑖_ values (**Fig. 3)**, and supported by LOOIC comparisons showing that models with this random phylogenetic effect outperform those without (e.g., model 8 vs. 1 or 43 vs. 10; **Table S9**; see **Table S13** for the group-level – i.e., random – effects by *Quercus* species). Second, closely related oaks also tend to host the same particular *Agrilus* species (*j*), an effect quantified by *α_2_*. Although smaller than the spatial effect *α_1_* (**Fig. 3** and **Fig. S2**), this signal is still important: models including *α_2_* outperform those excluding it (e.g., model 1 vs. 4 or 121 vs. 113; **Tables S9 and S11**). The best-performing metric for this effect is the minimum phylogenetic distance to another known host (𝑥*_2_*_𝑖𝑗_).

The phytophagy level of each *Agrilus* species – i.e., the number of oak species it exploits – is already encapsulated by the spatial and phylogenetic terms (*α_1_* and *α_2_*), since models that include an *Agrilus* species random effect for species-level prevalence perform no better (e.g., model 9 vs. 1 or 20 vs. 2; **Table S9**), or have very small effect sizes (**Table 1**, **Fig. S3**, **Table S3**). Similarly, including a proxy for sampling effort has little impact on explanatory power (e.g., model 7 vs. 1 or 88 vs. 48; **Tables S9 and S10**), or has very small effect sizes (**Table 1**, **Table S3**).

### Predicting *Agrilus* host status

Our chosen model predicts *Quercus*–*Agrilus* interactions effectively (**Fig. 4, Figs. S4 and S5**), with the log-odds scores for observed interactions being much higher (median log-odds = −1.35) than those for the rest (median log-odds = −8.05; **Fig. 4a**). To characterise the model’s performance, we divided this continuous distribution into predicted positive interactions (meaning that the *Agrilus* beetle *j* is predicted to be able to exploit the oak species *i*) and negative interactions (it is not), using our binary threshold (log-odds = −4.15; see **Materials and Methods**). Of the known interactions, 90.5% are correctly predicted to be positive; 90.6% of the interactions scored as 0s for model fitting are predicted to be negative (**Fig. 4b**). We hypothesise that the lower left-hand quadrant of **Fig. 4b** (predicted positive, but appear negative in the training dataset, i.e. “false positives”) includes *Quercus*–*Agrilus* interactions that either occur but have not yet been reported, or could occur in the future (these interactions include some recently recorded in *Agrilus* invasive ranges that were coded as negative for model training; see **Discussion – model performance**). Consequently, they identify potential hosts at risk (**Table S14**). This higher risk category includes a total of 696 interactions, encompassing all 32 *Agrilus* species and 105 *Quercus* species (including all 50 known hosts in our dataset).

**Fig. 4:**
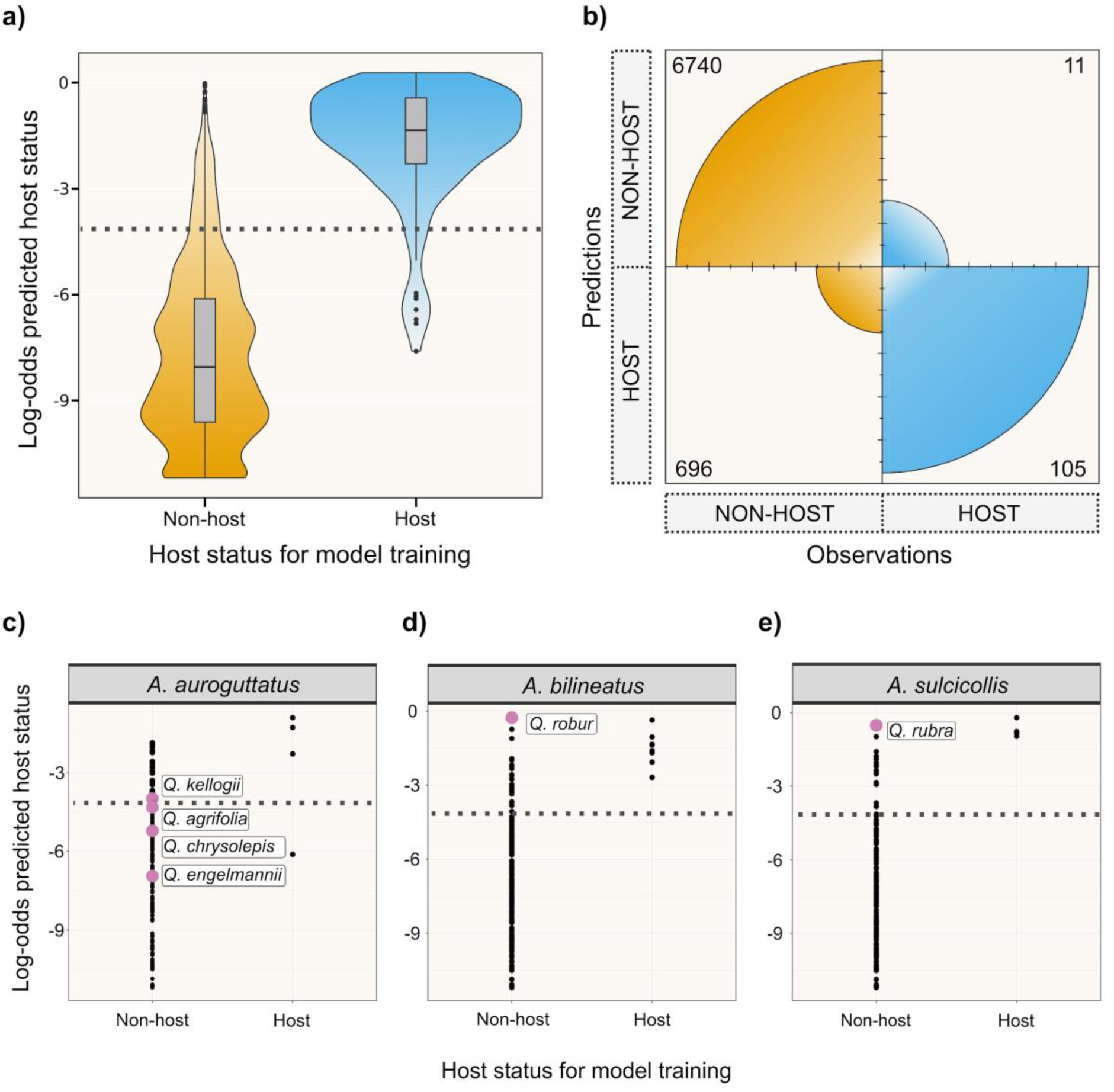
Continuous and binary predictions for the chosen model. **a) Continuous predictions (log-odds) of *Agrilus* host status for each *Quercus* species – *Agrilus* species pair.** The x-axis represents the observed host status for each *Quercus* species – *Agrilus* species pair, and the y-axis shows the log-odds of the model prediction, with higher values indicating a greater probability of a “positive” interaction. The dotted line indicates the threshold used in *b)* to divide predictions into predicted non-hosts and predicted hosts (see **Materials and Methods**). b) Binary predictions of *Agrilus* host status for each *Quercus* species – *Agrilus* species pair. Numbers indicate the number of *Quercus* species – *Agrilus* species pairs that belong to each quadrant. 90.52% of known *Quercus* species – *Agrilus* species interactions are predicted as such, and 90.64% of unobserved interactions are predicted as negatives. We hypothesise that the lower left-hand quadrant (696 *Quercus* species – *Agrilus* species pairs in total) might indicate future potential *Quercus*–*Agrilus* interactions and thus, potential future hosts at risk (**Table S14**). c-e) Continuous predictions for *A. auroguttatus*, *A. bilineatus* and *A. sulcicollis*, respectively. The dotted line represents the binary threshold. Pink labelled points indicate oak species that have been reported as new hosts in the introduced ranges of these beetles (scored as non-hosts for the purposes of our model).

To evaluate the model’s performance when information about a particular interaction is wrong or missing, we conducted two types of Leave-One-Out (LOO) analyses. In the first (**Fig. 5a**), known positive interactions were coded as 0s, assessing the ability of our model to correctly identify known hosts when they are incorrectly scored as non-hosts; this effectively simulates currently unrecognised host species. In the second analysis (**Fig. 5b**), missing interactions were incorporated using approximate Leave-One-Out predictions, estimating model predictions as if each interaction was missing from the training data.

**Fig. 5:**
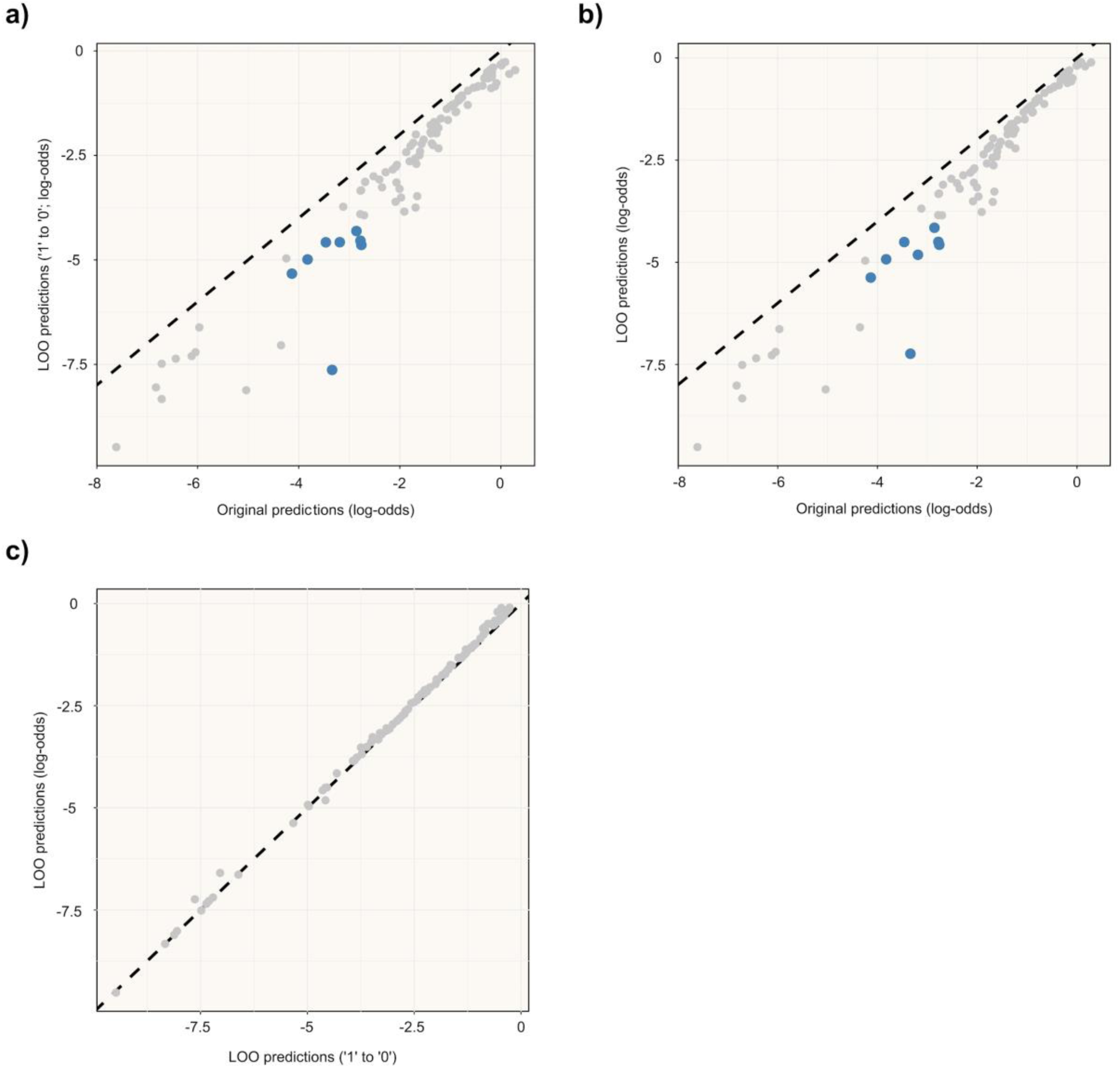
Leave-One-Out (LOO) predictions. Only shown for the 116 known interactions (i.e., *Quercus*–*Agrilus* pairs where the oak species is known to host the given beetle) in our dataset. **a) Predictions from the more stringent analysis where each 1 (i.e., known “positive” interaction) was converted into a 0 at a time, against the original predictions from the model. b) Predictions from the analysis where Leave-One-Out estimates were used to assess the effect of missing interactions, against the original predictions from the model. c) Predictions from the second LOO analysis (shown in *b)* against predictions from the first one (shown in *a)*).** The dashed grey line indicates the 1:1 line and the blue points indicate values that are over the binary threshold on one axis (i.e., in the “original predictions” in the case of *a)* and *b)*) but not the other; there are none in the case of *c)*, illustrating that these approaches serve as good proxies for each other.

Both analyses yield similar results, with individual log-odds prediction values showing a high degree of agreement (**Fig. 5c**). This indicates the second approach, which is less conservative and computationally intensive, would be sufficient for this assessment of model performance. In both cases, the model correctly classifies the oak as a potential host in 83.6% of instances, suggesting the model could be used to identify host species at risk from *Agrilus*, even where knowledge of existing host ranges is incomplete. Only eight interactions are predicted as positive in the original analysis but as 0 in the LOO analyses (**Fig. 5a, 5b**). These are: *Q. castaneifolia* – *A. graminis*, *Q. arizonica* – *A. parkeri*, *Q. prinoides* – *A. obsoletoguttatus*, *Q. ilicifolia* – *A. arcuatus*, *Q. durata* – *A. angelicus*, *Q. shumardii* – *A. quercicola*, *Q. marilandica* – *A. geminatus*, and *Q. acutissima* – *A. ussuricola*. Finally, we compared the second LOO approach against a five-fold cross-validation analysis (**Fig. S6**). Predictions were highly consistent, with strong Pearson correlations (r = 0.93 across all values and r = 0.97 for known positives), further supporting the robustness of our modelling approach to missing interactions.

## Discussion

In this study, we have investigated whether current patterns in the association between species of *Agrilus* beetles and *Quercus* trees can be used to predict future novel interactions, which have potential to be severely damaging to the newly acquired hosts. Such interactions have recently arisen, as in the case of the emerald ash borer (*A. planipennis*) (Herms & McCullough, 2014), which is decimating *Fraxinus* (ash) populations in North America, and *A. mali*, which is attacking apple trees in western China (Bozorov et al., 2019). These examples emphasise the need for effective approaches to predict the next major threat, so that mitigation can be put in place.

### The importance of phylogenetic and spatial variables in predicting future interactions

In the past, efforts to disentangle host–enemy dynamics have often integrated phylogenetic and trait data. For instance, Mech et al. (2019) devised a logistic regression model for predicting the future risk posed by non-native insects to North American conifers, and Barrow et al. (2019) employed host environmental, trait, and phylogenetic information to construct phylogenetic mixed-effects models (using brms, Bürkner, 2017) to predict parasite infection of birds. While both approaches successfully modelled host–enemy interactions, our model provides accurate predictions while maintaining simplicity, as it incorporates only three variables and does not rely on trait data, which can be arduous to compile (albeit some trait databases are available, e.g., Kattge et al., 2020). Instead, our mixed-effects model uses relatively simple predictors of host status, based on readily available phylogenetic and geographic information which are typically more centralised, accessible and generalisable.

In our model, geographic range overlap is a strong predictor of host use, supporting the idea that ecological opportunity plays a role in shaping host–pest associations. An oak species is more likely to host a given *Agrilus* species if at least part of its distribution overlaps with that of another host, as repeated exposure to a pest increases the likelihood of host jumps (Gilbert & Parker, 2016). This variable also likely captures other relevant patterns, such as beetle polyphagy; a generalist *Agrilus* species is more likely to be present on nearby trees, which may explain why the model performed well without a species-level random effect (see **Results – Variable predictors of *Agrilus* host status**). Range overlap may also reflect ecological similarities between plants with similar distributions, which can lead to convergent traits and shared climatic tolerances (Losos, 2011; Givnish, 2016; Wang et al., 2025), partly substituting for more explicit environmental niche modelling. This single predictor therefore encapsulates multiple ecological drivers of host use, allowing for a simple model that is both efficient and interpretable.

The two distinct phylogenetic effects captured in our model are consistent with the presence of a phylogenetic signal in the host range of *Agrilus* species that use oaks (see **Results – Phylogenetic signal in host status**). This evidence for phylogenetic signal aligns with that observed in other plant–enemy interactions (Parker & Gilbert, 2004; Gilbert & Webb, 2007; Pearse & Hipp, 2009; Yguel et al., 2011; Gilbert et al., 2012; Pearse & Altermatt, 2013; Barrow et al., 2019; Mech et al., 2019; Cirtwill et al., 2020; Pearse & Rosenheim, 2020; Lynch et al., 2021), including in *Quercus* herbivores (Pearse & Hipp, 2009), consistent with the idea that many host traits are phylogenetically conserved (Gilbert & Parker, 2016). Many of these studies also make use of phylogenetic information for predictive purposes. For example, Pearse & Altermatt (2013) used a simple generalised linear model to predict associations between German Lepidoptera and introduced plant species, using host breath information and minimum phylogenetic distance of each plant species to any known host of each lepidopteran. They built their model using information on native interactions and used it to predict values for non-native species, validating the results with known host status (Pearse & Altermatt, 2013). Both the presence and strength of this phylogenetic signal are, however, scale-dependent, with larger scales expected to result in stronger signals (Kamilar & Cooper, 2013; Pearse et al., 2013). For example, Cavender-Bares et al. (2006) proposed that the likelihood of observing phylogenetic clustering was higher in broadly-defined communities; they found that functional traits of plant communities in Florida (US) tended to be conserved at broad phylogenetic scales (woody plants), but failed to find this same pattern when focusing on a single lineage (*Quercus*) (Cavender-Bares et al., 2006).

In our model, the general phylogenetic tendency included as a random effect reflects a baseline preference of *Agrilus* to use closely related oak species, with a bias towards certain *Quercus* clades (**Fig. 2**). This variable might also capture ascertainment bias – as oak species which have attracted more observations seem to be non-randomly distributed through the phylogenetic tree (see host-rich clades in **Fig. 2**) – and may partly explain why our proxy for sampling effort made no discernible contribution to the model (see **Results – Variable predictors of *Agrilus* host status**). In contrast, the *Agrilus* species-specific phylogenetic signal makes the weakest contribution to model predictions, likely because the dataset is already constrained to *Agrilus* that use oaks. As such, the general random effect is probably capturing most of the phylogenetic information; if a *Quercus* species is a common host, other *Agrilus* species that exploit oaks might also be able to utilise it. Part of this effect might also be captured by the geographic metric (consistent with Tobler’s law; Tobler, 1970), as closely related species tend to occur nearer each other due to shared evolutionary history and habitat filtering (Gilbert & Parker, 2016). In fact, the oak genus itself is divided into geographically structured clades (Kremer & Hipp, 2020), meaning that more closely related hosts are likely to also be closer in space.

Previous studies have also sought to combine phylogenetic and geographic data when forecasting biotic interactions. For example, Robles-Fernández & Lira-Noriega (2017) used phylogenetic data to predict hosts of ambrosia beetles and subsequently looked at host distribution to highlight geographical hotspots of interactions, whereas Albery et al. (2020) built models based on phylogenetic proximity and geographic overlap, amongst other variables, to predict viral sharing between mammal hosts. However, to the best of our knowledge, our efforts represent the first attempt to integrate phylogenetic and geographic information to build simple and robust models for predicting future interactions within a plant–pest network. Given that phylogenetic data and occurrence records are already available for many taxa, together with a growing number of examples for phylogenetic signal in host ranges, we believe our approach could prove useful for predicting future interactions in many other systems.

### Model performance

Our model (**Fig. 4**) successfully classifies *Quercus*–*Agrilus* interactions with high accuracy, predicting 90.5% of known interactions, and achieving 83.6% accuracy in Leave-One-Out analyses. Notably, the lower left quadrant of **Fig. 4b** comprises 696 interactions not currently reported within the beetles’ native ranges. While these predictions represent false positives with respect to the input data, our validation analyses indicate the model can detect plausible interactions even when they are recorded as negative. As such, we believe this quadrant captures both overlooked and potential future interactions (**Table S14**).

Three *Agrilus* species in our dataset, i.e., *A. bilineatus*, *A. sulcicollis* and *A. auroguttatus*, were recently introduced to areas outside their native ranges, where they are have been observed exploiting novel *Quercus* hosts (Jendek & Grebennikov, 2009; Coleman & Seybold, 2011; Coleman, Graves, et al., 2012; Petrice & Haack, 2014; Haack & Petrice, 2019). These were treated as non-hosts in our analyses to test predictive capacity (**Fig. 4c–e**). The model successfully predicted 50% of these associations. *Agrilus bilineatus* and *A. sulcicollis* both have their recently recorded hosts (*Q. robur* and *Q. rubra*, respectively) ranked as the top novel prediction (**Fig. 4d–e**). *Agrilus auroguttatus* shows a more complex pattern; only one of four reported novel hosts (*Q. kelloggii*) exceeds the binary threshold, while another (*Q. agrifolia*) falls just below it (**Fig. 4c**). This reflects the fact that, while all four novel hosts are geographically distant from the known hosts of this beetle, *Q. kelloggii* and *Q. agrifolia* (both within host-rich clades, **Fig. 2**) are phylogenetically more closely related to other *Agrilus* hosts (**Fig. S7**). Interestingly, *A. auroguttatus* is the only one of these three beetles with a known host (*Q. peduncularis*) under the prediction threshold (**Fig. 4c**). In total, only ten *Agrilus* species have a known interaction under this threshold (**Fig. 4b**), typically involving single-host species or phylogenetically and geographically isolated oaks (**Fig. S7**). This suggests the model may be less reliable in cases of unusual host use patterns, cryptic diversity, or taxonomic uncertainty (e.g., *A. auroguttatus* was historically synonymised with *A. coxalis*; Hespenheide, 1979; Coleman, Lopez, et al., 2012).

The geographic distribution of the predicted hosts largely mirrors the overall distribution of *Quercus*, and that of known *Quercus* hosts (**Fig. S8)**. This is logical, as these regions provide a greater number of species that might act as hosts and more opportunity for novel interactions to arise (see **Fig. S8c-f** for maps showing the number of known and predicted hosts per region). Phylogenetically, predictions broadly (but not entirely) reflect known host-rich clades (**Fig. S9**). This suggests that, while detecting host-rich clades is a helpful first step, this approach will miss some vulnerable species.

### Implications of model predictions

Individual *Quercus* and *Agrilus* species involved in many predicted novel interactions may be of particular interest when considering where best to focus efforts to mitigate future risks. Two oak species, *Q. acutissima* (with two known interactions, **Table S1**) and *Q. rubra* (with six), stand out as potentially suitable hosts for all 32 *Agrilus* species. While our model might be overestimating their hosting ability, these species seem strong candidates for general hosts, with wide distributions (when considering both native and introduced ranges; GBIF, 2022) – and thus geographically close to other hosts – and falling within host-rich phylogenetic clades. Nevertheless, *Agrilus* species may vary in their ability to exploit them. For example, introduced species lacking natural enemies and escaping host defences (Brockerhoff & Liebhold, 2017) might be more aggressive and able to take over the niche (Work & McCullough, 2000; Fortuna et al., 2022). Alternatively, *Agrilus* species already established might take precedence over non-aggressive newcomers. Nevertheless, co-occurrence of multiple *Agrilus* species on a single host may be enabled by preferences for attacking different parts of trees (e.g., while most *Agrilus* larvae tend to girdle through the trunk, *A. angelicus* prefers young twigs; Swiecki & Bernhardt, 2006), or trees at different stages (e.g., *A. hastulifer* predominantly utilises dead trees, DEFRA, 2025, and *A. horni* prefers young suckers, Nord et al., 1965). In addition to *Q. acutissima* and *Q. rubra*, a further 26 *Quercus* species are predicted hosts for ten or more different species of *Agrilus* (**Table S14**). Likewise, there are some *Agrilus* species that are involved in a very large number of predicted novel interactions. For example, *A. obsoletoguttatus* is predicted to have 52 novel potential oak hosts. Interestingly, with ten reported host species and nine different host genera (Jendek & Poláková, 2014), it is, in fact, a generalist beetle capable of using a diverse range of hosts. While this species has not been reported causing harm to hosts in the past, it has also not yet escaped its native range (GBIF, 2025). Two further *Agrilus* species are predicted to have more than 40 novel oak hosts – *A. graminis* and *A. angustulus* (**Table S14**). In common with *A. obsoletoguttatus*, both species have numerous known hosts, including multiple oak species (nine in the case of *A. graminis* and ten for *A. angustulus*; **Table S1**), and thus their polyphagous nature is already documented.

### Limitations of the approach

We believe our model provides a robust framework for predicting future interactions between *Agrilus* beetles and their host plants, which can be extended to other enemy–host systems. However, some limitations and considerations must be acknowledged. Perhaps most obviously, it relies on the availability of phylogenetic information for the host taxa, which may not always be present. However, as more comprehensive datasets become accessible (e.g., Zuntini et al., 2024), use of phylogenetically-informed models, such as ours, is increasingly feasible. In addition, while the Leave-One-Out cross-validation analysis demonstrates our model can infer unreported interactions, systematic geographic biases in host reporting (Jendek & Poláková, 2014) may still influence the results. As mentioned above, we expect the bias towards some phylogenetic clades (**Fig. 2**) to partly reflect ascertainment bias. Furthermore, while the overall distribution of oak species broadly agrees with that of hosts (**Fig. S8**), China – where many oak species are present but few hosts predicted – is a notable exception, suggesting our model may be underpredicting interactions for this region.

Whether a species actually becomes a host will depend on factors additional to those accounted for in our model. The probability of a host jump will depend on ecological factors such as plant relative abundance (Gilbert & Parker, 2016) – which influences the likelihood of repeated encounters –, abiotic plant stress (Chamorro et al., 2015) – which can alter the individual tree susceptibility, e.g., (Showalter et al., 2018) –, niche availability and resource competition (Gidoin et al., 2015), or community structure (Gougherty & Davies, 2021). Pest spread can be facilitated in communities with closely related hosts (Gougherty & Davies, 2021), although polyphagous and monophagous pests may respond differently to habitat heterogeneity (with the former preferring species-rich habitats) (Brockerhoff et al., 2006). *Agrilus* species with different diet breadths could therefore thrive in different environments, even if they share some hosts. Still, most *Agrilus* species (ca. two-thirds) are only known to exploit one plant genus (Jendek & Poláková, 2014), so may perform better in homogeneous habitats. In fact, Dutch elm disease led to increased *Fraxinus* (ash) dominance in eastern North American forests (Gandhi & Herms, 2010; Barnes, 2011), which are now suffering from the *A. planipennis* (emerald ash borer) invasion (Gandhi & Herms, 2010). Moreover, some potential hosts may only occur in areas with environmental conditions otherwise unsuitable for pest survival, preventing establishment. Finally, regions with high trade activity may face elevated risks of species introductions. In future work, trade information could be used to complement our model by identifying routes or areas at high risk, similarly to Montgomery et al. (2023), who integrated global trade connectivity, alongside other variables, into a stochastic network model to estimate the probability of pest and pathogen entry and establishment. In sum, we do not anticipate every predicted interaction (**Table S14**) will materialise, as we expect a gap between potential and realised host ranges (Braga & Janz, 2021). Rather, our approach is intended to prioritise likely interactions for further assessment (e.g., ground-truthing) before translating these results into policy actions.

Even where predicted interactions do materialise, not all potential hosts will be equally affected. There is evidence of a conserved signal in host vulnerability for plant–pest interactions (Gilbert et al., 2015), meaning that only a subset of host species will suffer severe damage. Unfortunately, for *Agrilus* and many other genera, data on the severity of impact is much less readily available than that of host status. Nonetheless, hosts that are not severely damaged themselves can still facilitate transmission to others. For example, alternative hosts have been reported to act as enemy reservoirs that increase the risk of transmission to host crops (Saeed et al., 2015; Ocimati et al., 2018).

Finally, hosts currently within the native ranges of *Agrilus* beetles that do not seem to be heavily impacted could suffer increased damage in the future. *Agrilus* beetles are known to cause greater damage during periods of abiotic stress to hosts (Wei et al., 2004; Haack & Petrice, 2019) and when in association with other biotic agents (Brown et al., 2015). Climate change may alter host distribution and ecosystem composition (Chen et al., 2011), potentially leading to associations with other biotic stressors or facilitating novel interactions (Lancaster, 2020). It may prompt the planting of exotic trees considered to be better adapted to future conditions (Holl & Brancalion, 2020) inadvertently bringing highly susceptible hosts into contact with native pests (Dang et al., 2022). In fact, it has been proposed that global warming might have already prompted the apparent expansion of *A. biguttatus* in the UK (Brown et al., 2015). All these processes could result in native beetles causing significant damage in their current range, which in turn may facilitate their spread and interaction with naïve hosts.

In this study, we have set out to assess the risk that *Agrilus* species pose to *Quercus* trees. We have detected a phylogenetic signal in the distribution of oak hosts of *Agilus* and built a relatively simple yet powerful model that successfully predicts host–beetle interactions by using readily available phylogenetic and geographic data. We hope these findings will be used to focus efforts and inform actions aimed at preventing future damaging interactions, and to aid the development of targeted approaches for early pest detection in this and other host–pest systems.

## Acknowledgements

This study was supported by the Royal Botanic Gardens, Kew and Queen Mary University of London. E.H-G was supported by a studentship funded by the UK Government Department for Environment Food & Rural Affairs, through the Future Proofing Plant Health programme (grant no. TH1_10). This research utilised Queen Mary’s Apocrita HPC facility, supported by QMUL Research-IT (http://doi.org/10.5281/zenodo.438045). We thank A. Rossberg, the Evolution Lab Chat group at Queen Mary University of London and Plant Health and Adaptation Team at the Royal Botanic Gardens, Kew for useful discussions.

## Author contributions

L.J.K and R.A.N conceived and oversaw the project. E.H-G compiled the datasets and performed all analyses, with guidance from R.A.N and L.J.K. E.H-G wrote the manuscript, with input from L.J.K and R.A.N.

## Data availability

All scripts used in this study, as well as all lightweight input and output files (i.e., text files and phylogenetic input data files) are hosted on the following GitHub repository: https://github.com/elvirahg/agrilus-oaks-predictions. The full model outputs (.RData) are available on Zenodo at: doi.org/10.5281/zenodo.19537500, doi.org/10.5281/zenodo.19538352, doi.org**/**10.5281/zenodo.19538573, and doi.org/10.5281/zenodo.19538839.

## Supporting information (brief legends)

Fig. S1: Binary threshold used to divide predictions into “negative” and “positive” interactions

Fig. S2: Comparison of the simplest best-fit model (model 43) against models with only geographic or phylogenetic information

Fig. S3: Histogram of the estimate of the intercept value for the random effect of each *Agrilus* species in Model 65

Fig. S4: Continuous predictions for model 43 divided by oak species

Fig. S5: Continuous predictions for model 43 divided by *Agrilus* species

Fig. S6: Comparison of five-fold cross-validation log-odds predictions against Leave-One-Out (LOO) cross-validation log-odds predictions

Fig. S7: Value of different parameters for multiple *Agrilus* species with at least one host not predicted as such

Fig. S8: Choropleth maps showing the number of *Quercus* species, known oak hosts of *Agrilus*, and predicted hosts per country

Fig. S8: Phylogenetic patterns of clustering of *Agrilus* hosts across the *Quercus* phylogeny, also showing the distribution of predicted hosts

Table S1: Larval host information for *Agrilus* species that utilise *Quercus* species

Table S2: LOOIC cross validation to assess the best-performing geographic distance metric

Table S3: Model statistics

Table S4: LOOIC cross validation values for models 1–44 and comparisons against model with lowest LOOIC

Table S5: LOOIC cross validation values for models 45–88 and comparisons against model with lowest LOOIC

Table S6: LOOIC cross validation values for models 89–132 and comparisons against model with lowest LOOIC

Table S7: LOOIC cross validation values for best-fit models from all groups and comparisons against model with lowest LOOIC

Table S8: LOOIC cross validation values for those best-fit models from all groups with only contributing variables and comparisons against model with lowest LOOIC

Table S9: LOOIC group 1 model comparisons

Table S10: LOOIC group 2 model comparisons

Table S11: LOOIC group 3 model comparisons

Table S12: LOOIC comparison of best-performing models

Table S13: Group-level (i.e., random) effect for each *Quercus* sp. (i.e., each level) in model 43

Table S14: Interactions for which the *Quercus* species is not currently a confirmed larval host, but its predicted value in our statistical model is over the binary threshold (i.e., is it a predicted host)

Notes S1: Formulation of the geographic distance metrics

Notes S2: Formulation of geographic distance models

Notes S3: Formulation of the phylogenetic distance metrics

