## Supplementary Figures and Notes for "Combined phylogenetic and geographic data can predict plant–pest interactions with high accuracy"

The following Supporting Information is available for this article:

**Fig. S1**: Binary threshold used to divide predictions into "negative" and "positive" interactions

**Fig. S2**: Comparison of the simplest best-fit model (model 43) against models with only geographic or phylogenetic information

**Notes S2:** Formulation of geographic distance models

**Notes S3:** Formulation of the phylogenetic distance metrics

**Fig. S1: Binary threshold used to divide predictions into "negative" and "positive" interactions.** This threshold was obtained by computing the point that maximises both sensitivity (i.e., true positive rate, in blue) and specificity (i.e., true negative rate, in orange).

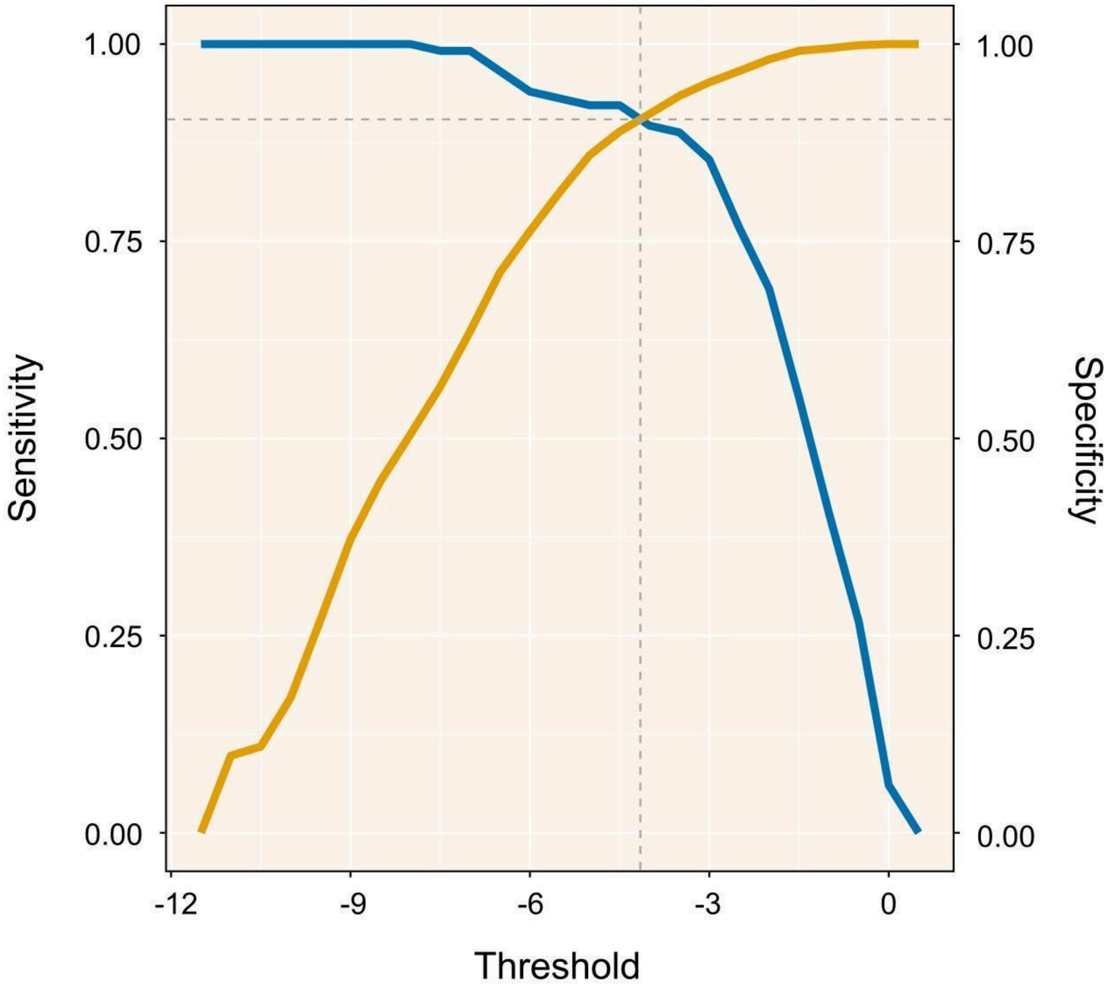

**Fig. S2: Comparison of the simplest best-fit model (model 43) against models with only geographic or phylogenetic information. a) Simplest best-fit model (model 43, y-axis) against a model with only geographic distance (model 2, x-axis) as an explanatory variable.** In the geographic-distance-only model, *Quercus* species with the same (normal-weighted) geographic distance to a host will have the same score values. **b) Model 43 against a model with only the phylogenetic distance fixed effect (model 4).** In model 4, *Quercus* species with the same phylogenetic distance to a host will have the same score values. **c) Model 43 against a model with only the phylogenetic clustering random effect (model 8).** In model 8, any given *Quercus* species will have the same values for each of the *Agrilus* interactions, and closely related oak species will have more similar values. **d) Model 43 against a model with only phylogenetic effects (both fixed and random, model 25).** Of note, the phylogenetic-only model (model 25) is more similar to the full one (model 43) than the geographic-only model (model 2), as shown by the $R^2$ values for these comparisons. **e) Model 43 against a model with geographic and phylogenetic (fixed) distance (model 10).** This shows the effect of adding the phylogenetic random effect to the model. **f) Model 43 against a model with geographic distance and the phylogenetic random effect (model 18).** This shows the effect of adding the phylogenetic distance (fixed) effect to the model, which is smaller than the rest (hence $R^2 = 0.98$). Dotted lines represent the linear relationship between the models (y = x). Blue points represent known positive *Agrilus* - *Quercus* host interactions.

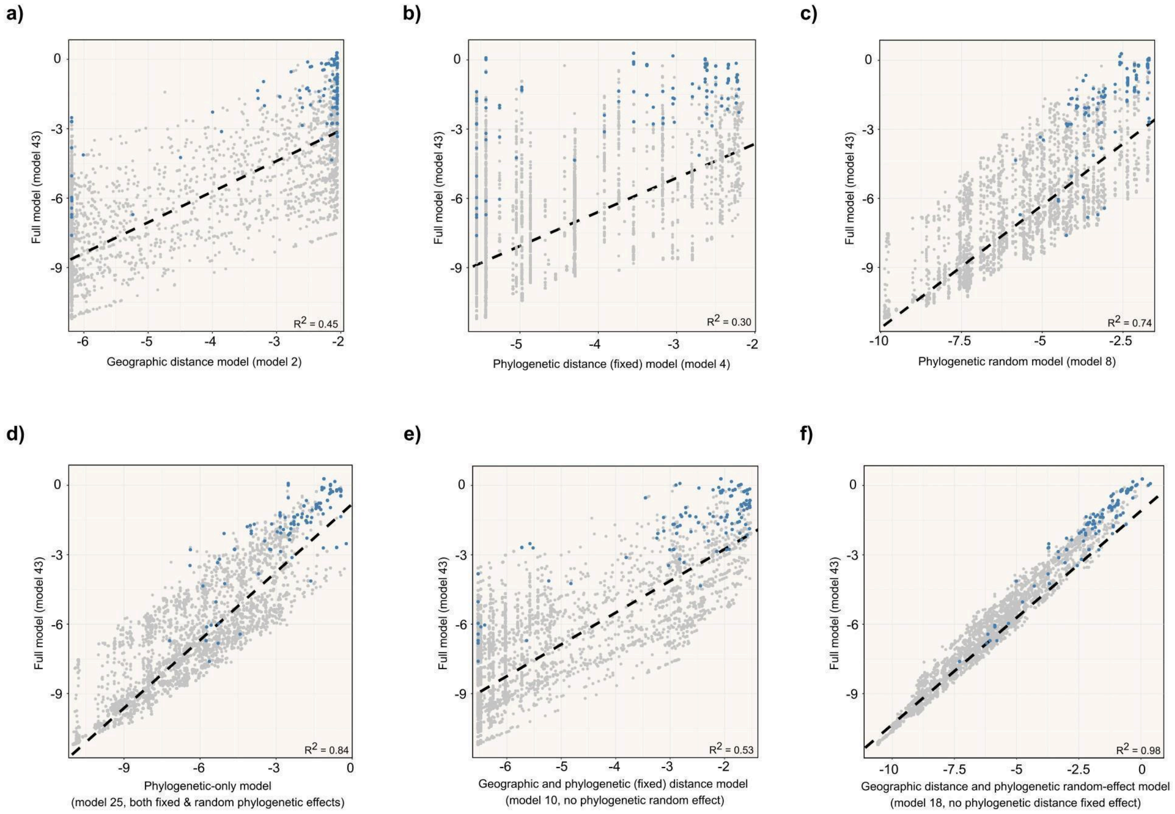
a)
Full model (model 43)
Geographic distance model (model 2)
R2 = 0.45
b)
Full model (model 43)
Phylogenetic distance (fixed) model (model 4)
R2 = 0.30
c)
Full model (model 43)
Phylogenetic random model (model 8)
R2 = 0.74
d)
Full model (model 43)
Phylogenetic-only model
(model 25, both fixed & random phylogenetic effects)
R2 = 0.84
e)
Full model (model 43)
Geographic and phylogenetic (fixed) distance model
(model 10, no phylogenetic random effect)
R2 = 0.53
f)
Full model (model 43)
Geographic distance and phylogenetic random-effect model
(model 18, no phylogenetic distance fixed effect)
R2 = 0.98

**Fig. S3: Histogram of the estimate of the intercept value for the random effect of each *Agrilus* species in Model 65.** This exemplifies the fact that the effect size of the random "*Agrilus* species" variable is small in all best-fitting models shown in **Table 1**.

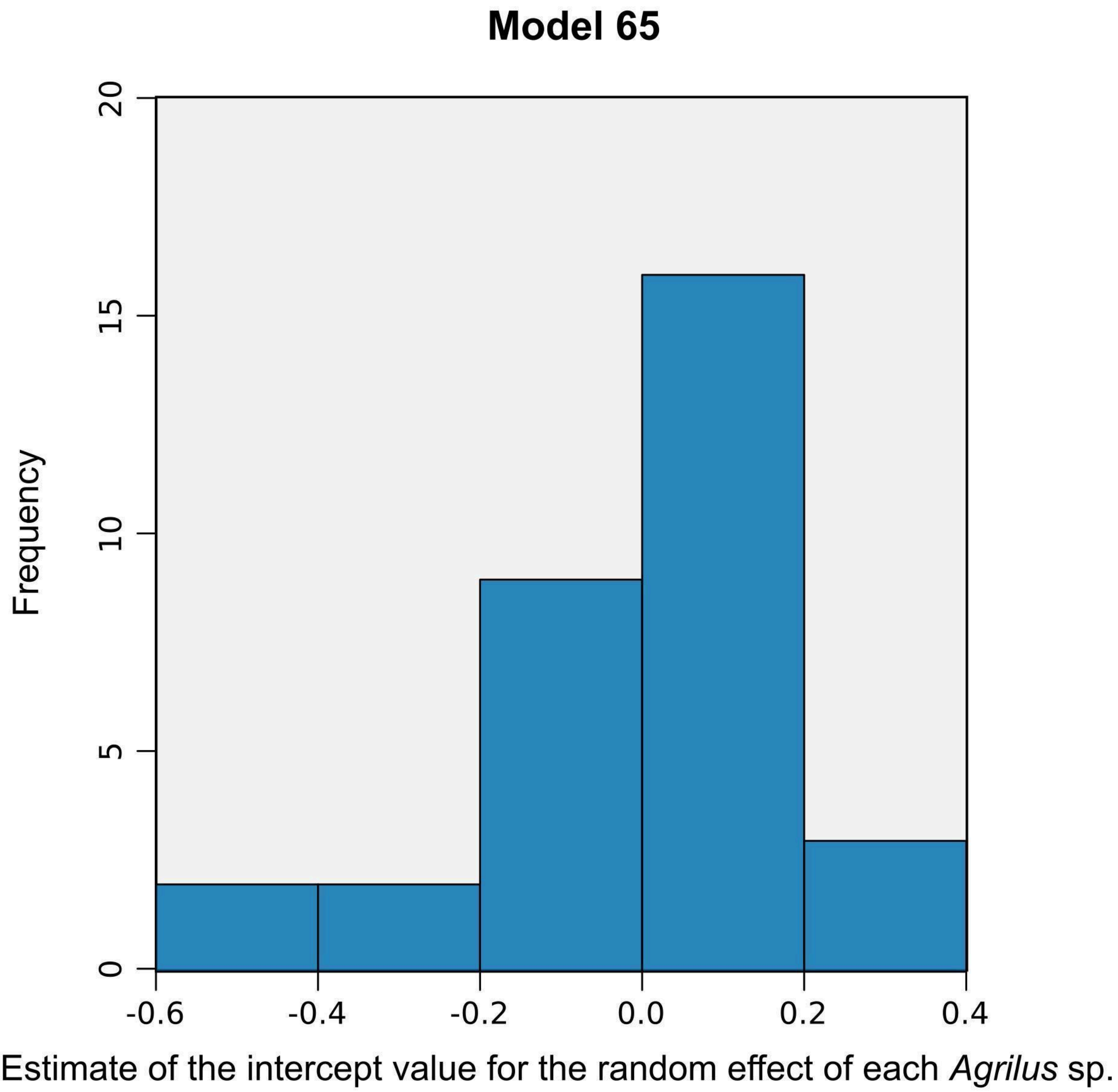

**Fig. S4: Continuous predictions for model 43 divided by oak species.** Each point represents an *Agrilus* species. The x-axis represents the observed interaction, with "0" if the given oak species is not known to host the given *Agrilus* species, and "1" if it is known to be host. The y-axis shows the log-odds of the predicted interaction for each *Quercus* species – *Agrilus* species pair, with higher values indicating a greater probability of a "positive" interaction. The dotted line indicates the binary threshold used to divide predictions into predicted non-hosts and predicted hosts.

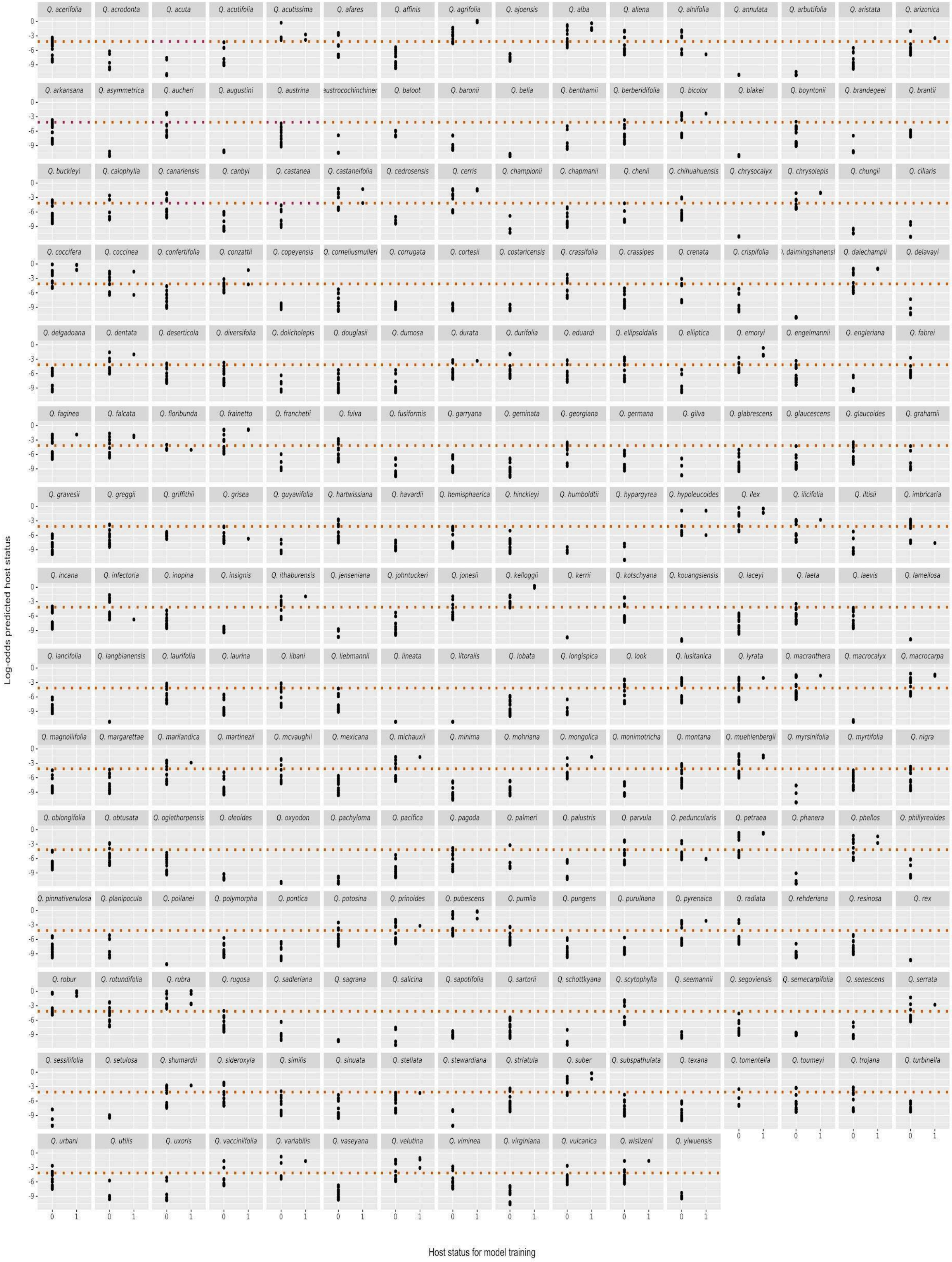
Log-odds predicted host status
Host status for model training

**Fig. S5: Continuous predictions for model 43 divided by *Agrilus* species.** Each point represents an oak species. The x-axis represents the observed interaction, with "0" if the given *Agrilus* species is not known to be hosted by the given *Agrilus* species, and "1" if it is known to be hosted by it. The y-axis shows the log-odds of the predicted interaction for each *Quercus* species – *Agrilus* species pair, with higher values indicating a greater probability of a "positive" interaction. The dotted line indicates the binary threshold used to divide predictions into predicted non-hosts and predicted hosts.

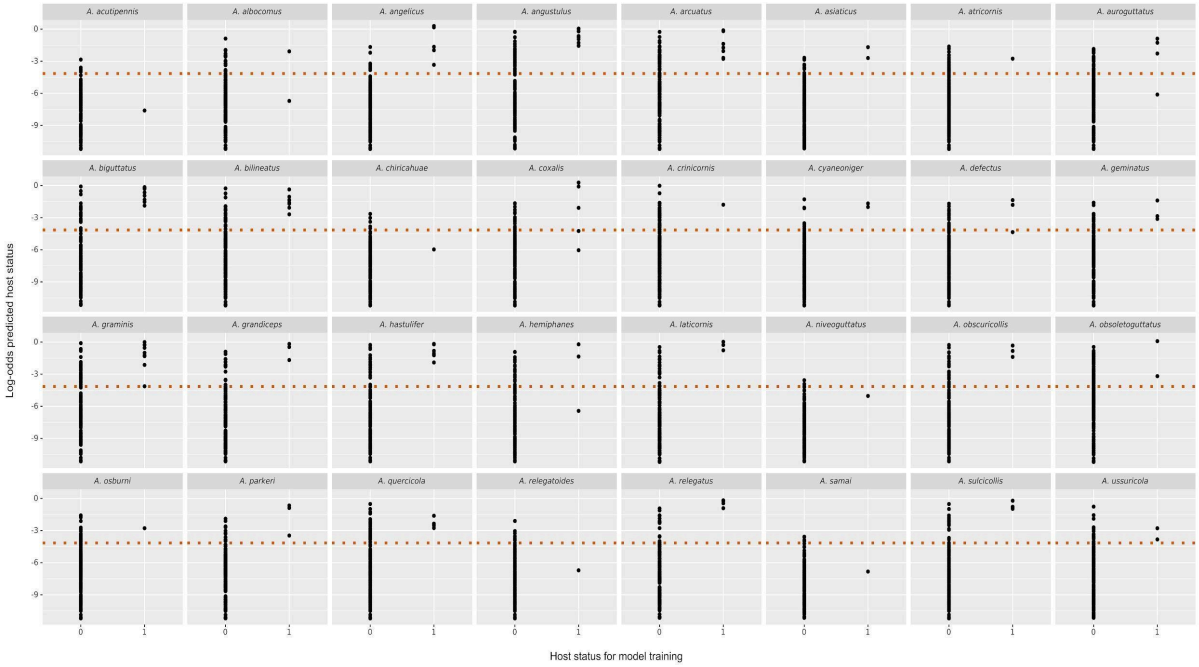

**Fig. S6: Comparison of five-fold cross-validation log-odds predictions against Leave-One-Out (LOO) cross-validation log-odds predictions.** Pearson's r = 0.93 across all values and r = 0.97 for known positives, indicating strong agreement between cross-validation approaches. While the accuracy of the five-fold analysis is higher (87.9% of known positive interactions correctly predicted vs. 83.7%), this analysis is also less generally conservative, yielding a higher number of total predictions above the binary threshold (14.75% vs. 10.70%). Interestingly, however, a higher number of originally predicted positive interactions (from the full model) are classified as negative under the five-fold approach (20 vs. 8). Blue represents known positive *Agrilus–Quercus* host interactions. Triangles indicate interactions predicted as positive (i.e. above the binary threshold). Horizontal and vertical dotted lines indicate the binary threshold (−4.15), and the diagonal represents the linear relationship between the models (y = x).

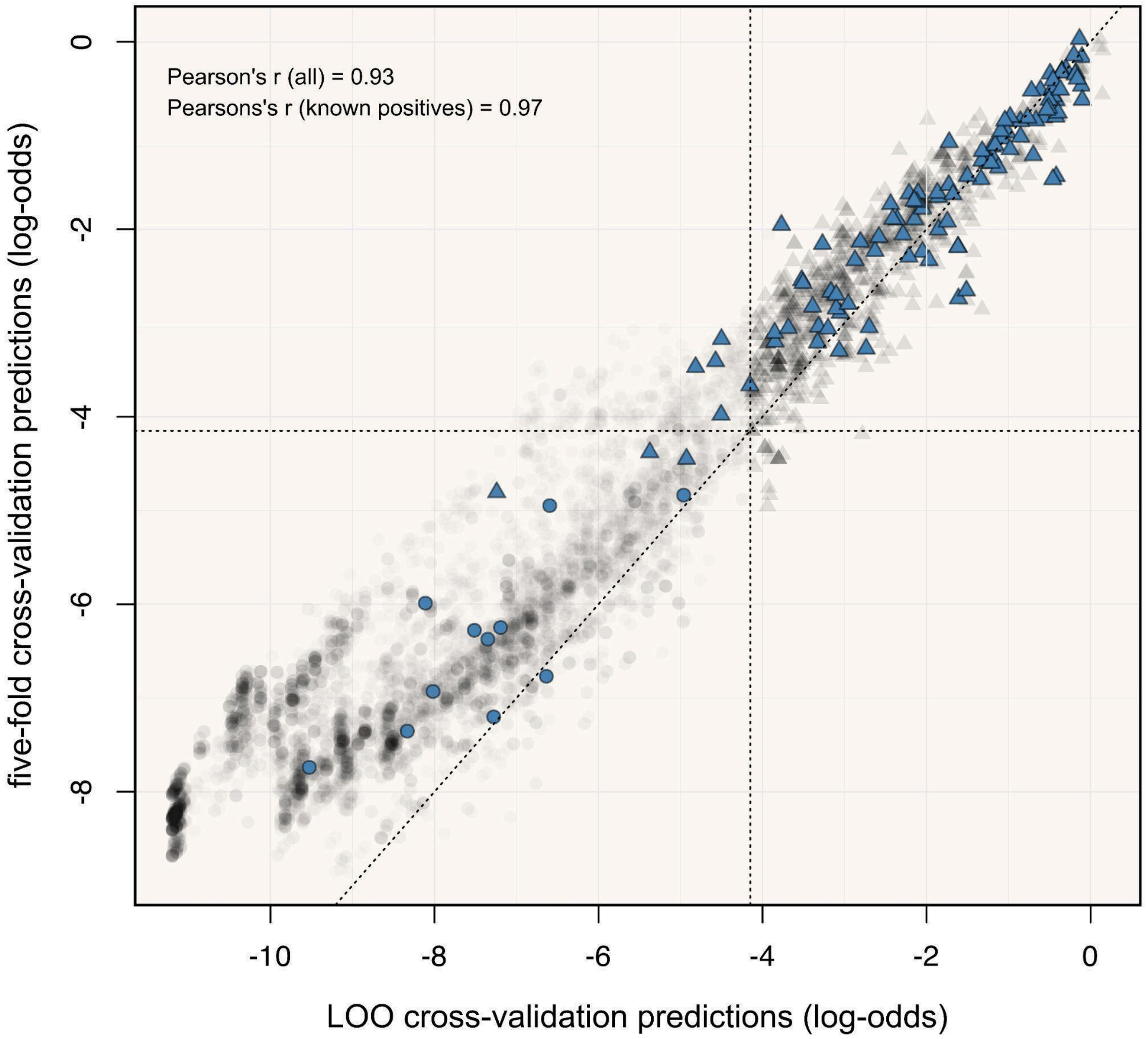

**Fig. S7: Value of different parameters for multiple *Agrilus* species with at least one host not predicted as such.** Blue indicates that the oak species is a known host of the given beetle in its current range, orange indicates that it is a novel host in its invasive range and pink indicates that it is a known non-host in the beetle's introduced range. Dotted lines indicate that the predicted value for the species fell under the binary threshold. **a, d, g, j, m) Value of the normalised geographic distance metric for each oak species to (other) known native hosts** (see **Materials and Methods**). **b, e, h, k, n) Minimum phylogenetic distance to any (other) known native host. c, f, i, l, o) Estimates of the phylogenetic random effect, which groups together oak species that are phylogenetically close.** This means that oak species closely related to known hosts of *Agrilus* beetles will have a higher chance of being predicted as a host in our model.

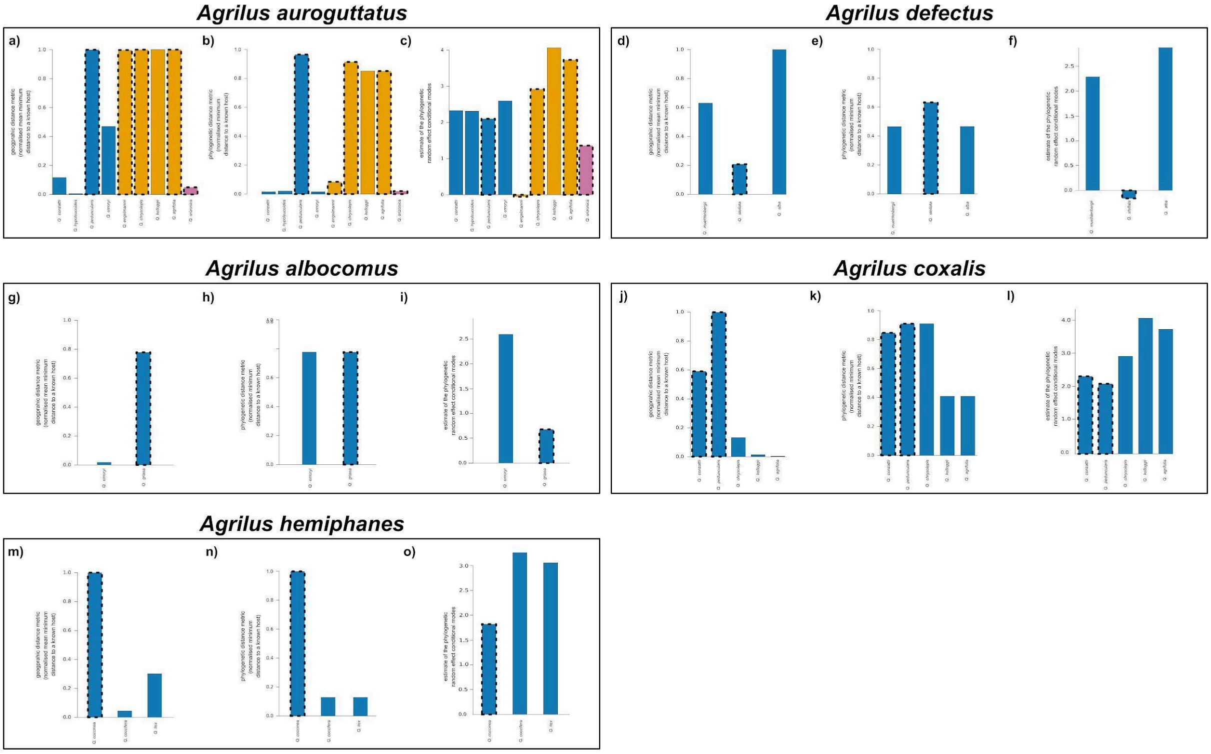

**Fig. S8: Choropleth maps showing the number of *Quercus* species, known oak hosts of *Agrilus*, and predicted hosts per country. a, c, e) Full distribution of native and non-native *Quercus* species b, d, f) Distribution of *Quercus* species in their native ranges.** Country level occurrences and native status were based on WCVP v10 data, retrieved through the R rWCVP package v1.3.0. Map lines delineate study areas and do not necessarily depict accepted national boundaries.

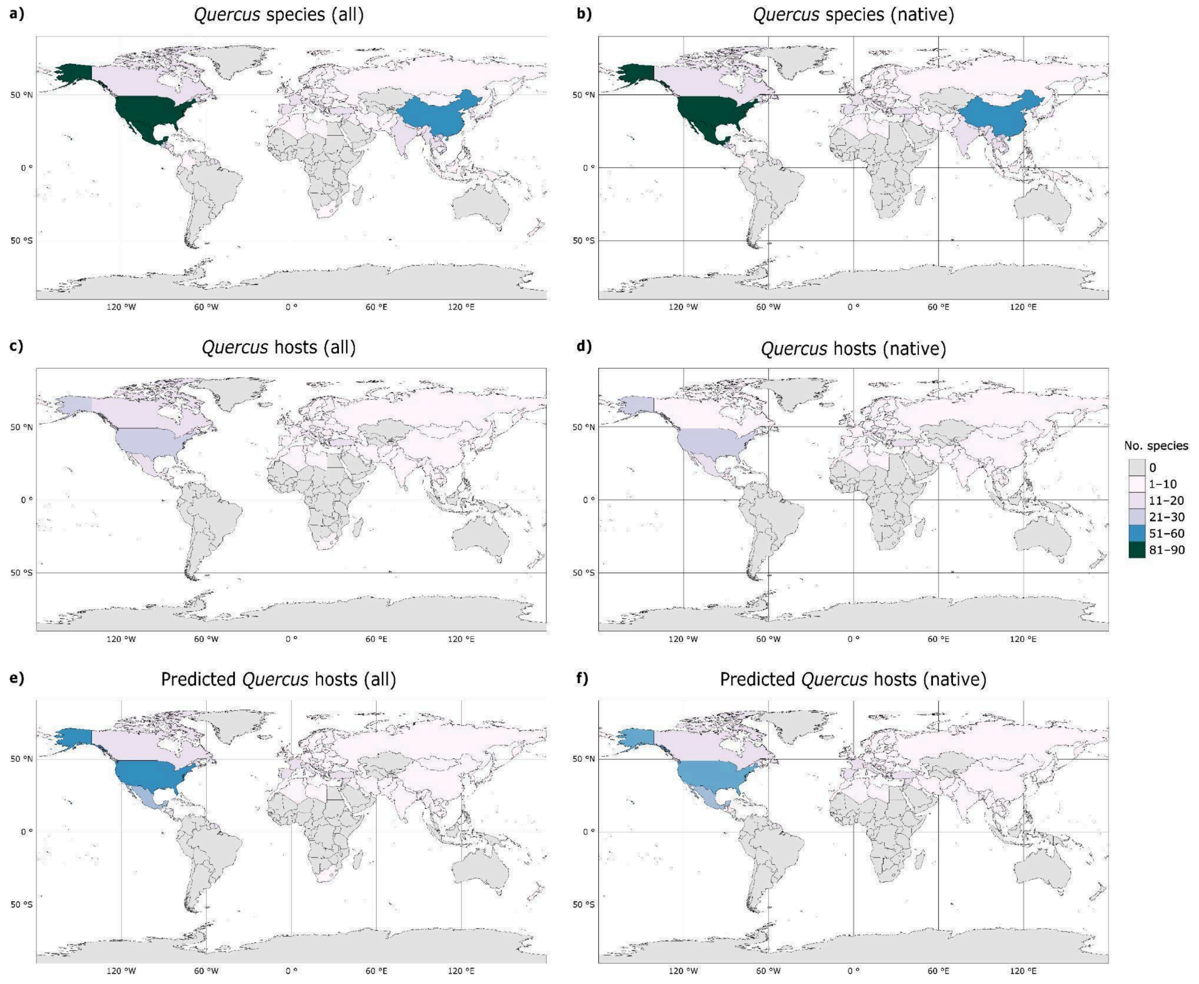

**Fig. S9: Phylogenetic patterns of clustering of *Agrilus* hosts across the *Quercus* phylogeny, also showing the distribution of predicted hosts.** Oak species labelled in orange are predicted as suitable hosts of at least one *Agrilus* species according to our model. Red and blue dots indicate that these nodes were identified as having significantly more or less descendants that are hosts than expected by chance, respectively. Grey clades indicate no over- or under-representation of hosts. Dark grey barplots indicate the number of known *Agrilus* species hosted per *Quercus* tree, and light grey circles indicate the number of cleaned GBIF entries (see **Materials and Methods**) available for each oak species. Stars indicate geographic region according to (Kremer & Hipp, 2020), and numbers indicate sections (see key). It should be noted that this clustering analysis is scale dependent.

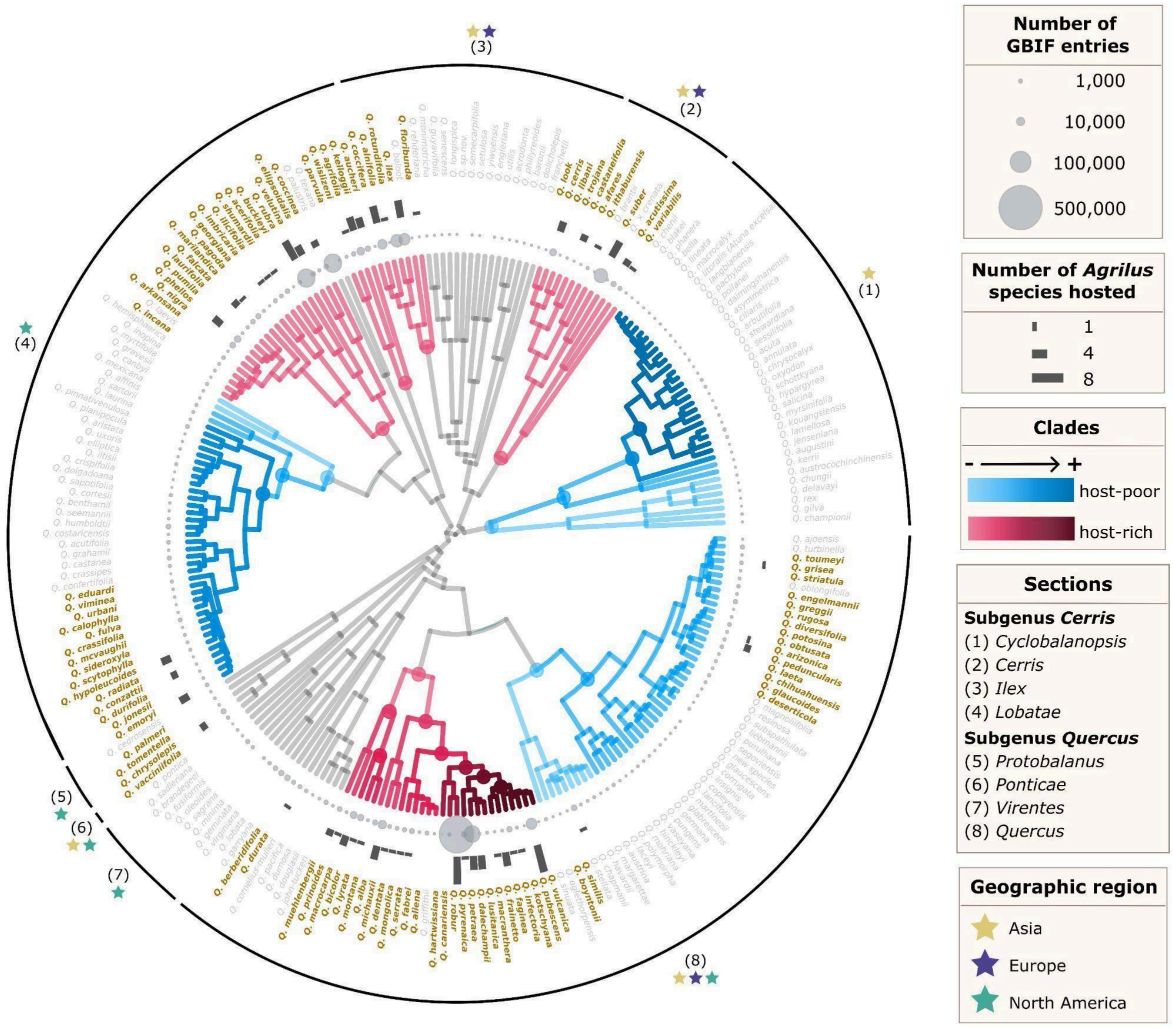

**Notes S1: Formulation of the geographic distance metrics**

We evaluated six metrics of spatial overlap between oak species and *Agrilus* hosts, calculated from geolocation information:

i) Mean of the minimum distances (in km) between each record of the given oak species and any known plant host of the given *Agrilus* species:

$$m_i = \frac{1}{|R_i|} \sum_{r \in R_i} min_{s \in S_i}\, d(r, s)$$

ii) Median of the minimum distances (in km) between each record of the given oak species and any known plant host of the given *Agrilus* species:

$$\tilde{m}_i = med_{r \in R_i}\left(min_{s \in S_i}\, d(r, s)\right)$$

Where $R_i$ is the set of occurrence records for oak species $i$, and $r$ is one specific record. For each species $i$, let

$$S_i = \bigcup_{h \in H,\, h \neq i} R_h$$

denote the set of occurrence records for all host species excluding those belonging to $i$, with $H_i$ being the set of host species associated with the focal *Agrilus* species (excluding $i$), and $R_h$ being the set of occurrence records for host species $h$. The function $d(r, s)$ denotes the geographic distance (in km) between occurrence records $r$ and $s$. If $i$ is the only known host species, the metric was set to 30,000 km.

iii) The transformed mean minimum distance to the corresponding normal probability density (μ = 0 km, σ = 50 km), multiplied by -1 (so that species close in space had lower scores) and rescaled to range from 0 to 1:

$$M_{i(norm)} = rescale(-\, \phi_{0,50}(m_i))$$

iv) The transformed median minimum distance to the corresponding normal probability density (μ = 0 km, σ = 50 km), also multiplied by -1 and rescaled to range from 0 to 1:

$$\tilde{M}_{i(norm)} = rescale(-\,\phi_{0,50}(\tilde{m}_i))$$

where $\phi_{\mu,\sigma}(x)$ is the normal probability density function with mean μ and standard deviation σ.

v) The log-transformed mean minimum distance, rescaled to range from 0 to 1:

$$M_{i(log)} = rescale(ln(m_i + 0.1))$$

vi) The log-transformed median minimum distance, rescaled to range from 0 to 1:

$$\tilde{M}_{i(log)} = rescale(ln(\tilde{m}_i + 0.1))$$

where a constant of 0.1 was added to avoid undefined values at zero distance.

**Notes S2: Formulation of geographic distance models**

We built separate models using each of the six geographic distance metrics, as defined in the following equation:

$$y_{ij} \sim Bernoulli(p_{ij})$$

$$logit_{p_{ij}} = \alpha_0 + \alpha_1 x_{1ij} + e_{ij}$$

Where $y_{ij} \in \{0, 1\}$ indicates if *Quercus* species $i$ can host the *Agrilus* species $j$. The probability $p_{ij}$ is a function of the following variables. The value $\alpha_0$ is the intercept, $x_{1ij}$ is the geographic distance metric from the *Quercus* species $i$ to any (other) known host of *Agrilus* species $j$, and $\alpha_1$ is the associated coefficient, and $e_{ij}$ is the normally distributed residual error.

**Notes S3: Formulation of the phylogenetic distance metrics**

We computed three phylogenetic metrics for each *Quercus–Agrilus* interaction using the time-calibrated *Quercus* phylogenetic tree:

i) the mean of the phylogenetic distance of the given oak species to (other) *Quercus* hosts of the given *Agrilus* beetle:

$$\frac{1}{|H\{i\}|-1} \sum_{h \in H,\, h \neq i} d(i, j)$$

ii) the minimum phylogenetic distance of the given oak species to any (other) *Quercus* hosts of the given *Agrilus* beetle:

$$d(i, j)$$

iii) the sum of both:

$$\frac{1}{|H\{i\}|-1} \sum_{h \in H,\, h \neq i} d(i, j) \; + \; d(i, j)$$

Where $H$ is the set of host oak species for a given *Agrilus* species, $d(i, j)$ is the phylogenetic distance between species $i$and $j$, and $d_{root} = 0.5$. If $i$ is the only known host, the metric was set to $2 \times d_{root}$.
